# The RNA Atlas, a single nucleotide resolution map of the human transcriptome

**DOI:** 10.1101/807529

**Authors:** Lucia Lorenzi, Hua-Sheng Chiu, Francisco Avila Cobos, Stephen Gross, Pieter-Jan Volders, Robrecht Cannoodt, Justine Nuytens, Katrien Vanderheyden, Jasper Anckaert, Steve Lefever, Tine Goovaerts, Thomas Birkballe Hansen, Scott Kuersten, Nele Nijs, Tom Taghon, Karim Vermaelen, Ken R. Bracke, Yvan Saeys, Tim De Meyer, Nandan Deshpande, Govardhan Anande, Ting-Wen Chen, Marc R. Wilkins, Ashwin Unnikrishnan, Katleen De Preter, Jørgen Kjems, Jan Koster, Gary P. Schroth, Jo Vandesompele, Pavel Sumazin, Pieter Mestdagh

**Affiliations:** Center for Medical Genetics, Ghent University, Belgium; Cancer Research Institute Ghent (CRIG), Ghent, Belgium; Texas Children’s Cancer Center, Baylor College of Medicine, Houston, USA; Illumina, San Diego, California, USA; Data Mining and Modelling for Biomedicine group, VIB Center for Inflammation Research, Ghent, Belgium; Department of Applied Mathematics, Computer Science, and Statistics, Ghent University, Ghent, Belgium; Department of Data Analysis and Mathematical Modelling, Ghent University, Belgium; Department of Molecular Biology and Genetics, Aarhus University, Denmark; Biogazelle, Zwijnaarde, Belgium; Department of Diagnostic Sciences, Ghent University, Belgium; Department of Pulmonary Medicine, Ghent University, Belgium; Department of Respiratory Medicine, Ghent University, Belgium; University Systems Biology Initiative, School of Biotechnology and Biomolecular Sciences, UNSW Sydney, Sydney, NSW, 2052, Australia; Adult Cancer Program, Lowy Cancer Research Centre, UNSW Sydney, NSW 2052, Australia; Prince of Wales Clinical School, UNSW Sydney, NSW 2052, Australia; Institute of Bioinformatics and Systems Biology, National Chiao Tung University, Taiwan; Academisch Medisch Centrum (AMC), University of Amsterdam, The Netherlands

## Abstract

The human transcriptome consists of various RNA biotypes including multiple types of non-coding RNAs (ncRNAs). Current ncRNA compendia remain incomplete partially because they are almost exclusively derived from the interrogation of small- and polyadenylated RNAs. Here, we present a more comprehensive atlas of the human transcriptome that is derived from matching polyA-, total-, and small-RNA profiles of a heterogeneous collection of nearly 300 human tissues and cell lines. We report on thousands of novel RNA species across all major RNA biotypes, including a hitherto poorly-cataloged class of non-polyadenylated single-exon long non-coding RNAs. In addition, we exploit intron abundance estimates from total RNA-sequencing to test and verify functional regulation by novel non-coding RNAs. Our study represents a substantial expansion of the current catalogue of human ncRNAs and their regulatory interactions. All data, analyses, and results are available in the R2 web portal and serve as a basis to further explore RNA biology and function.

## Main

The introduction and constant improvement of RNA-sequencing technologies have enabled us to interrogate the human transcriptome at nucleotide resolution, exposing distinct RNA biotypes beyond protein-coding messenger RNAs (mRNAs). A plethora of regulatory non-coding RNAs, including microRNAs (miRNA), long intergenic non-coding RNAs (lincRNAs), antisense RNAs (asRNAs) and circular RNAs (circRNAs) have been identified and are actively explored as novel players in human development and disease including cancer^1–3^. Several consortium-based efforts have contributed to the discovery and quantification of these RNA biotypes in heterogeneous sample collections^4–12^. The resulting transcriptome landscapes are serving as crucial community resources to study RNA biology, mechanism, function, and biomarker potential^13–16^. However, these studies have mostly applied RNA-sequencing technologies that profile the small- and polyadenylated-RNA transcriptomes. As a result, a systematic survey of the non-polyadenylated transcriptome and the circularized transcriptome, and their relationship to other RNA biotypes, is currently missing. To capture a more complete diversity of the human transcriptome, we applied three complementary RNA-seq methods on a heterogeneous collection of 300 human samples, including 45 tissues, 162 cell types, and 93 cell lines (Figure 1A, Supplemental Table 1). From these samples, we generated strand-specific small RNA (298 samples), polyA (295 samples) and total RNA (296 samples) libraries that were sequenced at a median depth of 13 M, 60 M paired-end, and 125 M paired-end reads respectively, resulting in a total of 125 Billion reads (Figure S1). By integrating these datasets, we assembled transcripts representing five major RNA biotypes, including mRNAs, lincRNAs, asRNAs, circRNAs and miRNAs, culminating in a stringently selected transcriptome and its matching expression atlas. Broad intron-coverage from the total RNA-sequencing data enabled data-driven prediction of transcriptional and post-transcriptional modulation of gene expression by non-coding RNAs, all of which are available through the R2 web portal.

**Figure 1.**
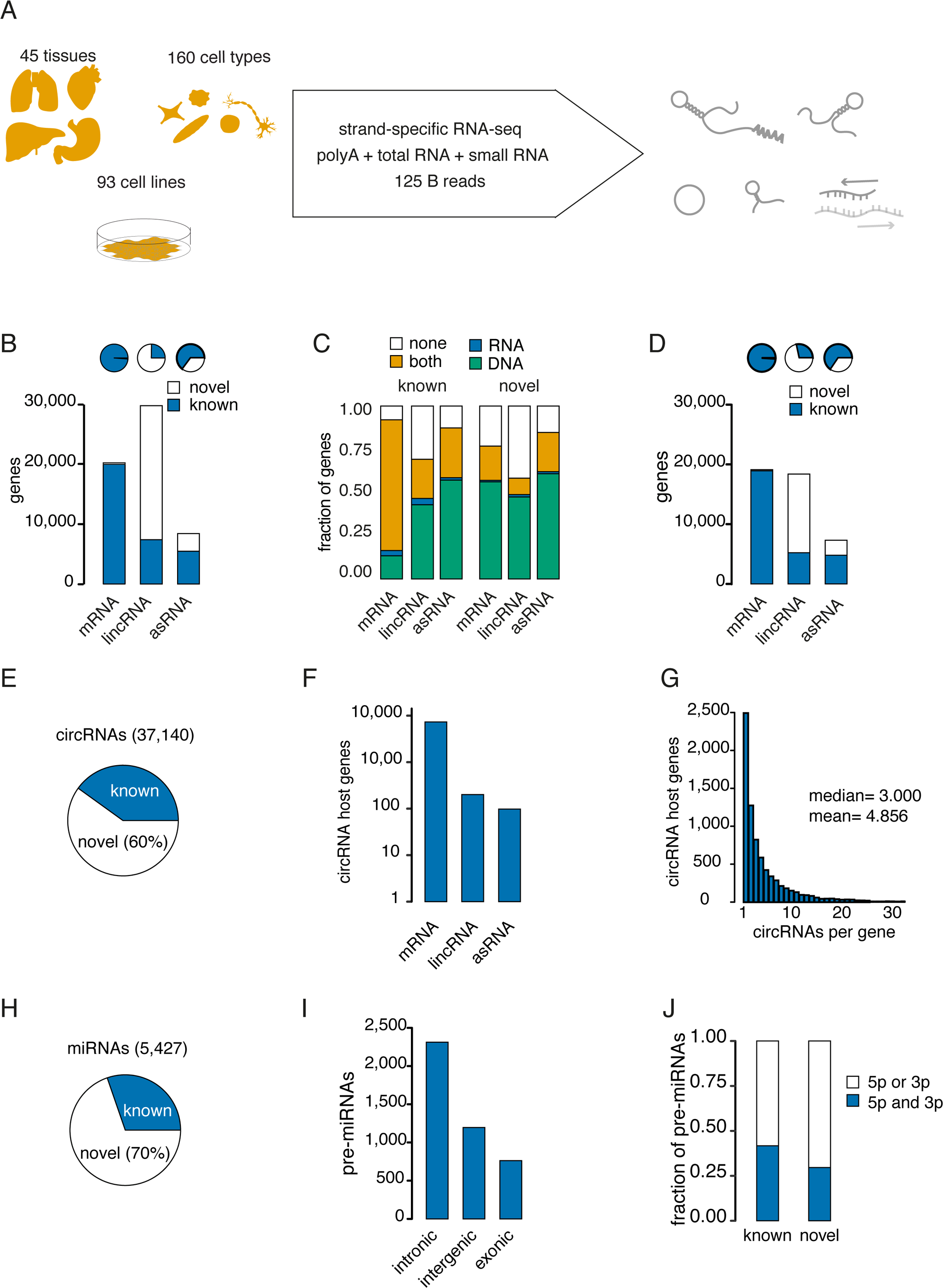
RNA Atlas transcriptome generation and annotation. (A) A heterogeneous collection of tissues, cell types and cell lines was sequenced through complementary strand-specific RNA-sequencing methods to profile the major RNA-biotypes in the human transcriptome. (B) Number and fraction of known and novel mRNA, lincRNA and asRNA genes in the complete RNA Atlas transcriptome. (C) Fraction of known and novel genes with independent evidence for transcriptional activity at the DNA level (chromatin states, green), RNA level (CAGE-peak, blue) and both levels (orange) or genes without independent evidence (white). (D) Number and fraction of known and novel mRNA, lincRNA and asRNA genes in the stringent RNA Atlas transcriptome. (E) Fraction of known and novel circRNA isoforms produced from genes in the stringent RNA Atlas transcriptome. (F) Number of stringent mRNA, lincRNA and asRNA host genes with expressed circRNA isoforms. (G) Histogram of expressed circRNA isoforms per host gene. (H) Fraction of known and novel candidate mature miRNAs. (I) Annotation of miRNA precursors based on location relative to genes in the stringent RNA Atlas transcriptome. (J) Fraction of known and novel pre-miRNAs with expressed mature miRNAs from both arms (blue) or one arm (white) of the precursor.

## Results

### Assembling a comprehensive human transcriptome reveals numerous single-exon lincRNAs

The basis of the RNA Atlas transcriptome was created by newly assembled mRNAs, lincRNAs, and asRNAs after rigorous filtering and merging with known transcripts from Ensembl annotation (v86)^17^ (Figure S2, Figure 1B, Supplemental Table 2, see Methods for details). To gather independent evidence supporting the transcriptional activity of these genes, we integrated public cap analysis of gene expression (CAGE) sequencing data (FANTOM^6^) and various chromatin states associated with transcription or with enhancer activity from comprehensive cell and tissue collections (Epigenomics Roadmap^18^). The majority of known (88%) and novel (62%) genes were closely associated—within 500 base pairs of transcription start sites (TSSs)—with a CAGE-peak or relevant chromatin state (Figure 1C). Non-coding RNA genes, and novel genes in general, were mainly supported by chromatin states while known mRNAs were supported by both CAGE-peaks and chromatin states. Genes supported by a CAGE-peak or chromatin state were retained in what we refer to as the stringent RNA Atlas transcriptome (Figure 1D). An additional 754 genes without CAGE-peak or chromatin state support, but with supra-median expression level (details in Methods), were also included. This transcriptome consists of 19,107 mRNA genes (of which 188 (0.9%) are novel), 18,387 lincRNAs (of which 13,175 (71%) are novel) and 7,309 asRNAs (of which 2,519 (34%) are novel). From the total RNA-sequencing data, we subsequently identified 38,023 circRNAs with more than 4 back-spliced reads (Supplemental Table 3). Of these, 37,140 were derived from stringent RNA Atlas genes and thus retained in the stringent RNA Atlas transcriptome. The majority (60%) of the circRNAs did not match previously annotated species in circBase^19^ and almost all (98%) were processed from mRNA host genes, with a median of 3 circRNAs per host gene (Figure 1E-G). In line with previous reports, circRNAs had a median number of 4 exons and were flanked by introns that were significantly longer than introns not flanking circRNAs (P < 1 x 10^−10^, Wilcoxon signed-rank test) (Figure S3). Finally, we identified 5,427 candidate mature miRNAs from the small RNA sequencing data, 70% of which were novel (Figure 1H, Supplemental Tables 4 and 5). Precursor transcripts for these miRNAs were predominantly located in intronic and intergenic regions and the fraction of precursors giving rise to both a 3p and 5p mature form was slightly lower for novel miRNAs (30%) compared to known (41%) miRNA genes (Figure 1 I-J). Precursor miRNAs are processed from larger primary miRNA transcripts whose transcription start sites are not easily defined (for instance, 1,196 precursors are located in intergenic regions). Therefore, we did not integrate CAGE-seq data or chromatin states to define a stringent set of miRNAs. Instead, we used expression-based relationships between the candidate mature miRNAs and their predicted targets to enrich for small RNA sequences that effectively show miRNA-like behavior. Further details on these analyses are presented in the penultimate section of the results (Figure 5).

**Figure 2.**
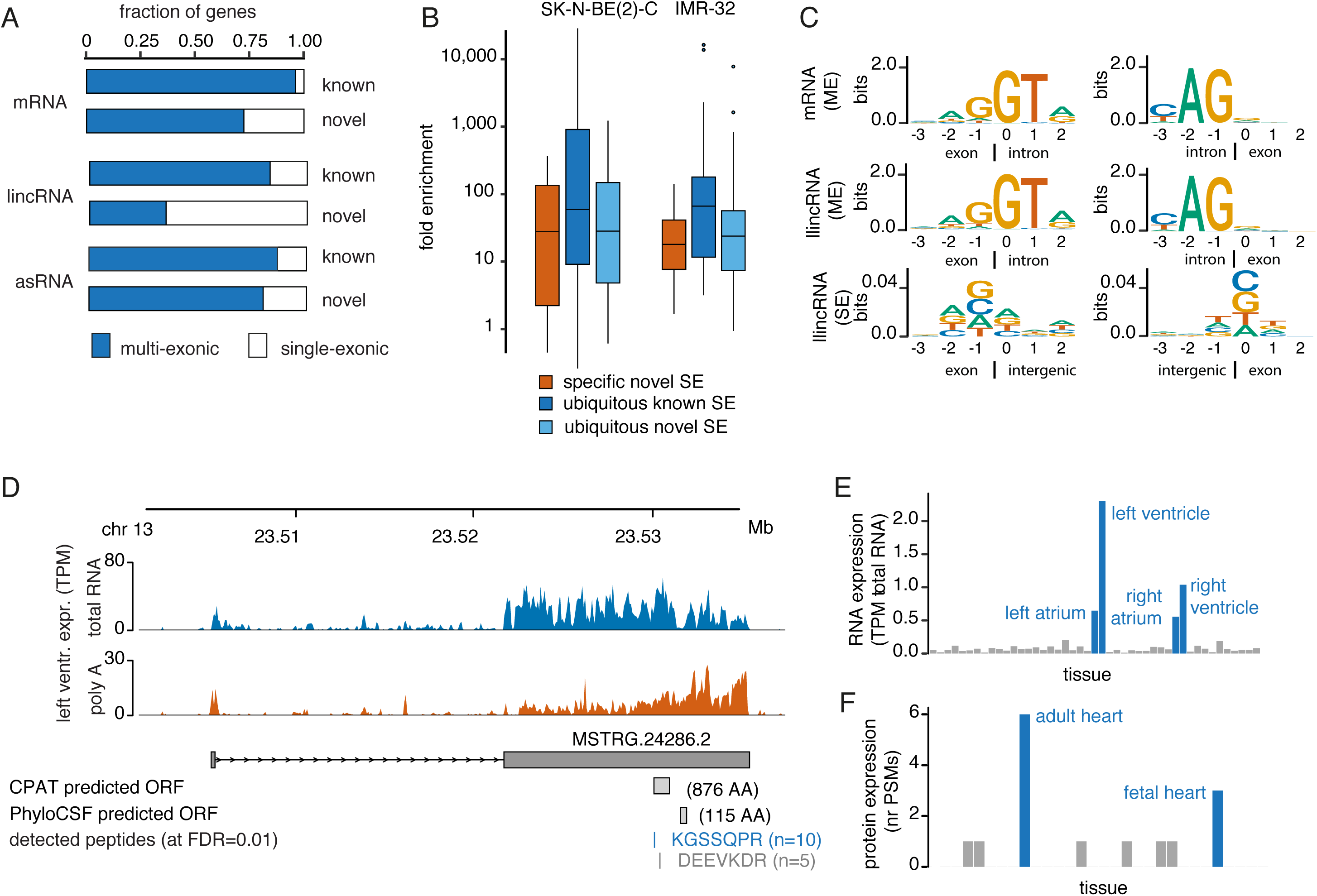
The RNA Atlas transcriptome reveals novel mRNAs and single-exon lincRNAs. (A) Fraction of single- and multi-exon genes in the mRNA, lincRNA and asRNA biotypes. (B) qPCR validation of novel single-exon lincRNAs in two RNA Atlas cell lines. The distributions of fold enrichment between RT and no-RT reactions are shown for ubiquitously expressed known and novel single-exon lincRNAs (dark blue, light blue, respectively) and for novel single-exon lincRNAs specifically expressed in each the cell line (orange). (C) Sequence motif analysis of the exon/intron boundary (exon/intergenic boundary for single-exon genes) for multi-exon mRNAs, multi-exon lincRNAs and single-exon lincRNAs. The y-axis shows the information content measured in bits. (D) Example of a novel heart-specific mRNA with matching peptides from public mass spectrometry data. Expression of this gene is specific for heart-tissue samples (blue bars) both at RNA (E) and peptide (F) level.

**Figure 3.**
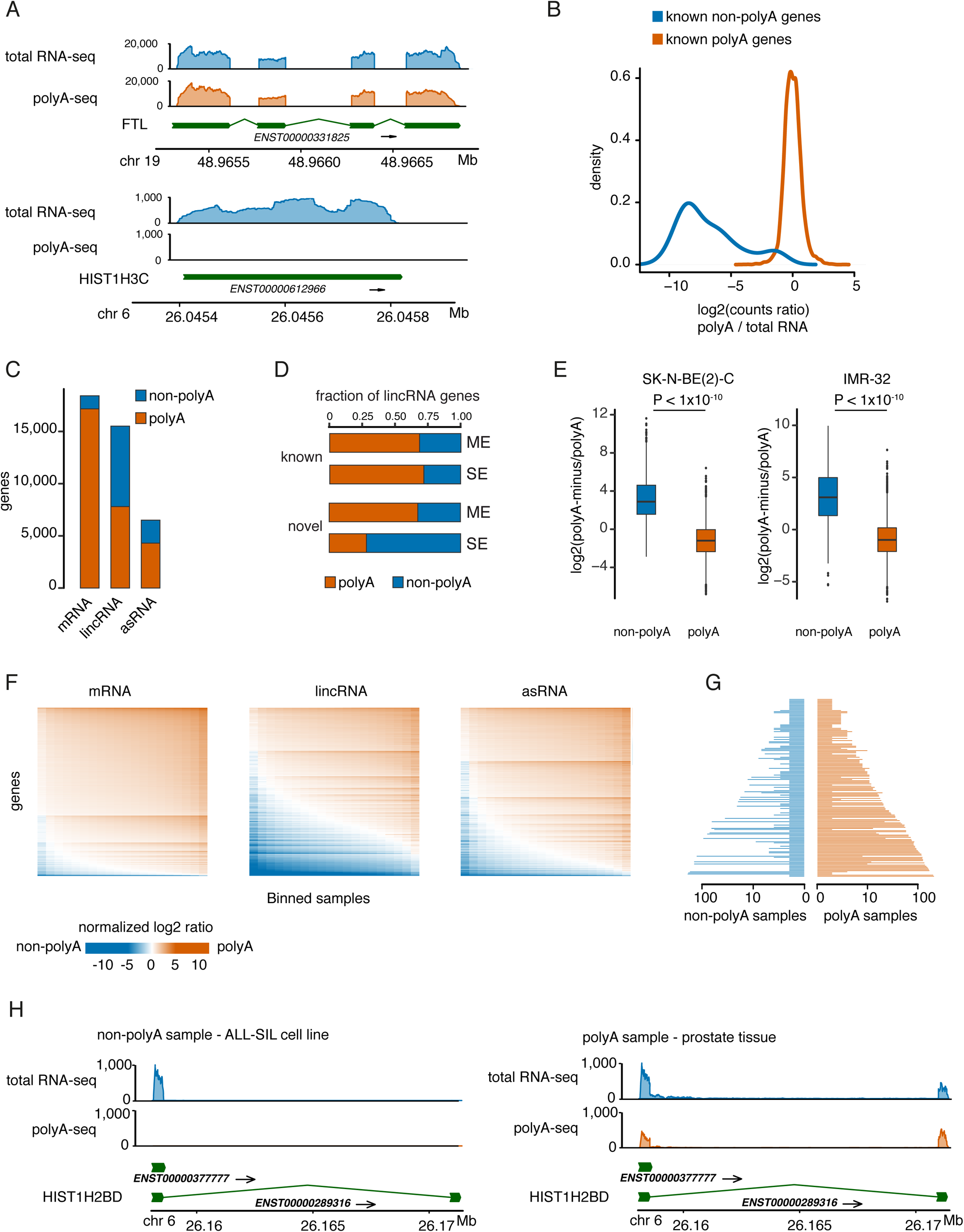
Analysis of polyadenylation status. (A) Read coverage profiles from polyA-sequencing and total RNA-sequencing libraries for a known polyadenylated gene (upper panel) and a known non-polyadenylated gene (lower panel). (B) Distribution of the normalized log2 ratio of counts from polyA-sequencing versus total RNA-sequencing in an individual sample (human umbilical vein endothelial cell) for known polyadenylated (orange) and non-polyadenylated genes (blue). (C) Number of polyadenylated and non-polyadenylated genes for the different RNA biotypes (majority vote across samples). Only genes in the stringent set, with 10 or more counts in at least one sample and with an uneven majority vote across samples are shown. (D) Fraction of polyadenylated and non-polyadenylated lincRNA genes (known and novel, single- and multi-exon). (E) Validation of non-polyadenylated genes through polyA-minus sequencing in two RNA Atlas cell lines. Boxplots show the log2 counts ratio between polyA-minus and polyA sequencing for genes classified as non-polyadenylated (blue) and polyadenylated (orange). (F) Heatmaps showing polyadenylation status of genes across samples. Only genes with 100 or more counts in at least 20 samples are shown. For the selected genes, all samples with at least 100 counts were sorted based on normalized log2 ratio and binned in a total of 20 bins (min samples per bin=1). For each bin, the mean normalized log2 ratio is plotted. (G) Selection of genes with varying polyadenylation across samples (see methods for definition of these genes). The number of samples in each category is shown, with values in log10 scale. Genes are sorted by evenness (i.e difference in number of samples in each category) and by number of polyadenylated samples (H) Example of a gene (HIST1H2BD) whose varying polyadenylation across samples can be explained by differential expression of alternatively polyadenylated isoforms. Coverage profiles from total RNA-sequencing and polyA-sequencing are shown for a non-polyadenylated sample (ALL-SIL cell line, left panel) and a polyadenylated sample (prostate tissue, right panel).

**Figure 4.**
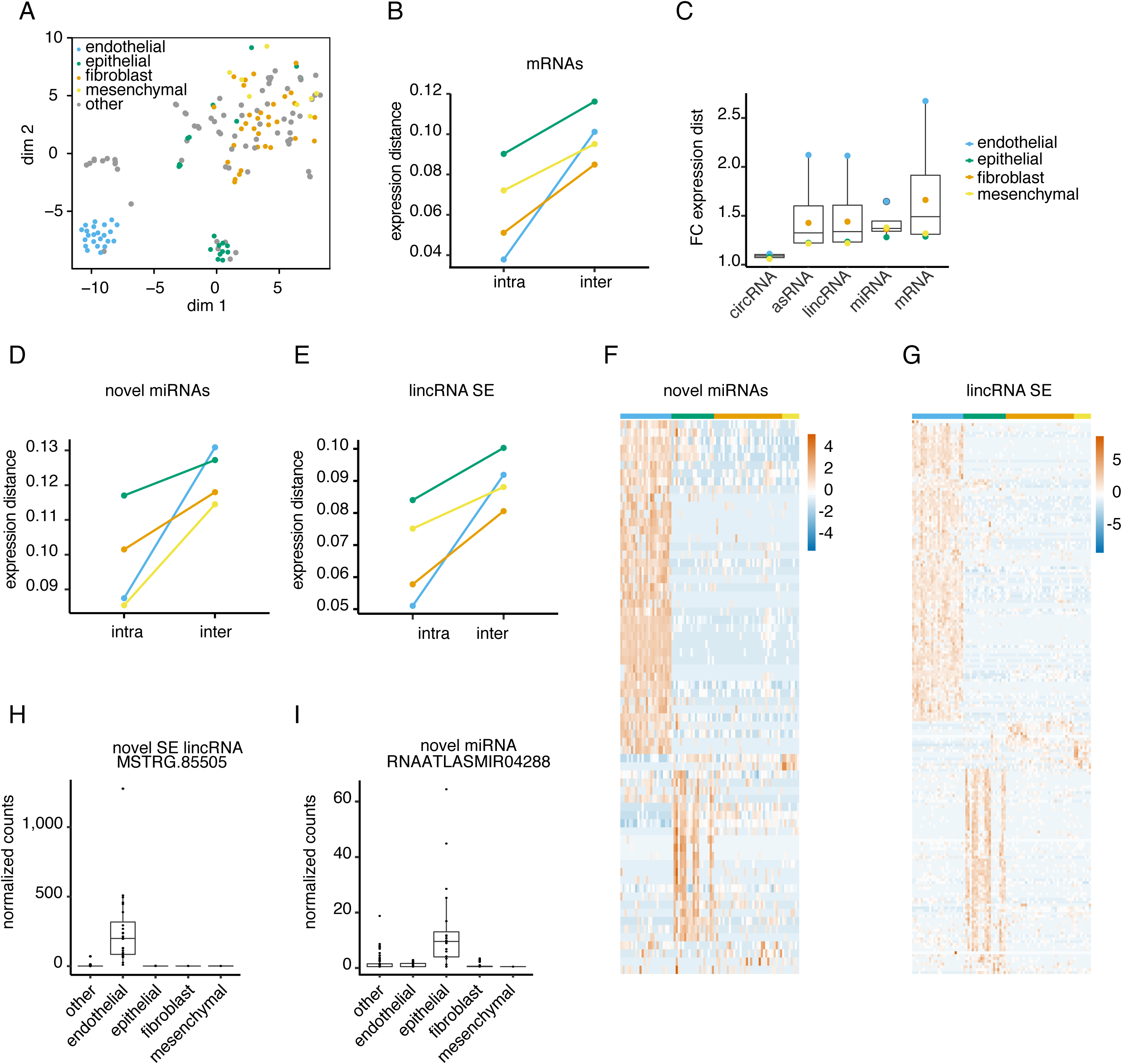
Association between sample ontology and expression-distance. (A) t-SNE plot of the RNA atlas cell types based on mRNA expression. Samples are colored according to the 4 biological cell subtypes of interest (epithelial, mesenchymal, fibroblast, endothelial). (B) For each biological cell subtype, the median mRNA expression-based distances between pairs of samples within a subtype (intra-distances) and between each sample from the subtype and all other samples (inter-distances) are shown. (C) Distribution of fold changes between median inter- and intra-distances calculated based on expression of the different RNA biotypes. (D) Median intra- and inter-distances based on expression of novel miRNAs. (E) Median intra- and inter-distances based on expression of single-exon lincRNAs. (F) Expression heatmap for novel miRNAs significantly upregulated in each of the cell subtypes. (G) Expression heatmap for single-exon lincRNAs significantly upregulated in each of the cell subtypes. (H) Example of an endothelial-specific single-exon lincRNA. (I) Example of an epithelial-specific novel miRNA.

**Figure 5.**
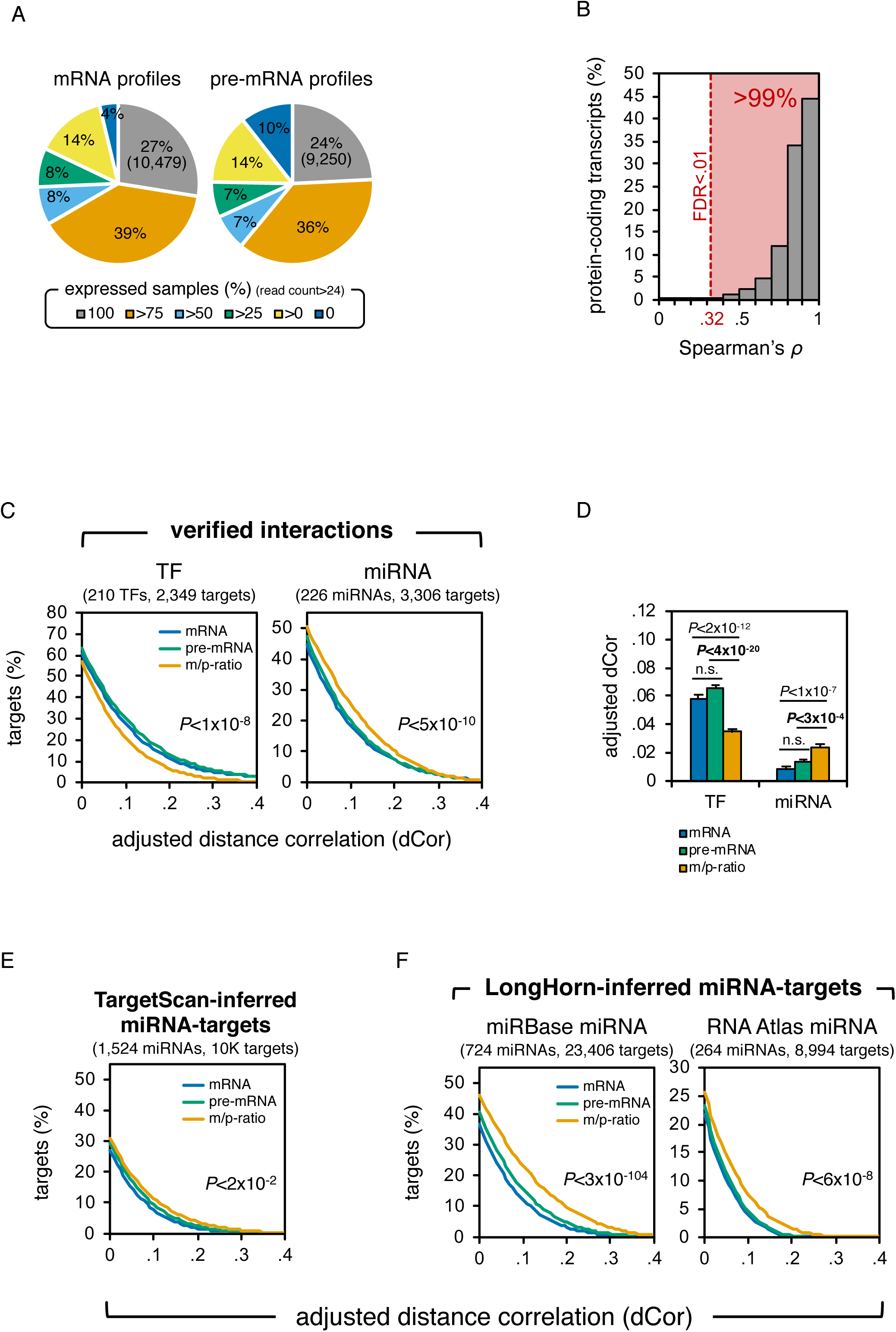
Total RNA transcriptomes facilitate the use of intron expression profiles to study regulatory modalities. (A) Distribution of protein-coding transcripts whose exonic (mRNA, left) and intronic (pre-mRNA, right) expression estimates are supported by at least 24 unique reads in 0, >0%, >25%, >50%, >75%, or 100% of RNA Atlas profiled samples. (B) Distribution of Spearman’s correlations between pre-mRNA and mRNA expression profiles (n=7,289, log2 transformed). (C) Adjusted distance correlation (dCor) between TF or miRNA expression profiles and their verified target’s pre-mRNA and mRNA expression profiles and the ratio between them (m/p-ratio); p-values are the geometric mean of the p-values for the differences between pre-mRNA - m/p-ratio and mRNA - m/p-ratio correlations. (D) Correlation with verified targets. On average, the adjusted distance correlations between the profiles of regulators and their verified target’s mRNA and pre-mRNA showed no significant difference, however, correlations between regulator and target m/p-ratio profiles were significantly lower and higher for TFs and miRNAs, respectively. (E) Sequence-based miRNA-target predictions by TargetScan did not exhibit improved miRNA-target m/p-ratio correlations. However, (F) LongHorn-inferred direct targets of miRBase and RNA Atlas-identified miRNAs did have improved regulator-target m/p-ratio correlations; p-values were estimated by the two-sample Kolmogorov-Smirnov test. The number of regulators and interactions tested is given in parentheses. Only miRNAs with at least 10 targets are shown.

Unsurprisingly, the majority of novel expressed loci in the stringent RNA Atlas transcriptome were annotated as lincRNAs. In contrast to mRNAs and asRNAs, most novel lincRNAs (65%) were single-exon genes (Figure 2A). Single-exon lincRNA genes are often removed from transcriptome assemblies as they are thought to derive from contaminating DNA. As we applied a stranded RNA-sequencing workflow, DNA fragments that get incorporated in the library would result in reads mapping to both strands in a nearly equal ratio. The mean exonic strandedness for single-exon lincRNAs was 96% and similar to that of multi-exon lincRNAs (94%) or mRNAs (95%), suggesting they do not originate from DNA fragments contaminating the RNA sample (Figure S4). This was further validated experimentally by qPCR for 110 single-exon lincRNA genes (48 novel ubiquitously expressed, 32 novel specifically expressed in one of 2 cell lines used for validation and 30 known) on RNA samples either or not reverse transcribed into cDNA. For those genes that were successfully amplified (Cq < 35, n=101), we observed a significant fold increase of qPCR signal in the RT versus no-RT samples (Wilcoxon signed-rank test, P < 1 x 10^−10^), further confirming that these single-exon lincRNAs originate from RNA and not residual DNA fragments (Figure 2B, Figure S5). The fact that all single-exon lincRNAs in the stringent RNA Atlas transcriptome have a CAGE peak (13%) or chromatin state(s) indicative for active transcription (98%) close to their TSS also favors this interpretation. Unlike multi-exon mRNAs or lincRNAs, and similar to single-exon mRNAs, single-exon lincRNAs are not flanked by canonical splice sites (Figure 2C, Figure S6 A). Moreover, the genomic distance between single-exon lincRNAs and the nearest up- or downstream exon is significantly larger (median 2.3 kb) compared to the genomic distance between consecutive exons in multi-exon mRNAs (median 0.511 kb, P < 1 x 10^−10^) or lincRNAs (median 1.17 kb, P < 1 x 10^−10^, Wilcoxon signed-rank test) (Figure S6 B). These observations support that the single-exon lincRNA genes are not part of multi-exon genes that we failed to assemble because of missing junction reads.

While most novel genes were identified among the ncRNA biotypes, our workflow also revealed a handful of novel candidate mRNAs. Based on in silico predictions, 188 novel genes showed high coding potential (Supplemental Table 6). The coding potential of these novel mRNAs was further substantiated through re-analysis of mass spectrometry data from the Human Proteome Map^20^, revealing peptides matching 33 (18%) of these novel mRNA genes (FDR < 0.01). We identified peptides whose expression profile across tissues correlated with that of their template mRNAs, underscoring the validity of the results. These include a novel heart-specific peptide whose template mRNA is specifically expressed in heart tissues (Figure 2D-F) and a peptide with high abundance in T-cells and B-cells whose template mRNA shows highest expression in spleen and lymph node (Figure S7). Protein domain analysis of the heart-specific protein sequence revealed a TLV_coat domain, also present in human syncytin genes. Homology to syncytin was further validated through BLAST analysis. Syncytin proteins mediate cell fusion during placental development, a process also occurring in muscle and cardiac tissue^21^.

### Integrating polyA and total RNA-seq data reveals thousands of non-polyadenylated genes

We then took advantage of our matching polyA and total RNA-sequencing data to establish the polyadenylation status of the stringent RNA Atlas genes. We reasoned that, after scaling count data from both libraries for differences in coverage, polyadenylated genes should show equal coverage in both polyA and total RNA libraries while coverage for non-polyadenylated genes should be low or absent in polyA libraries (Figure 3A). Indeed, the distribution of log2 count ratios (polyA/total RNA) for known polyadenylated genes is centered around zero and significantly higher than that of known non-polyadenylated genes (Figure 3B, Figure S8, Supplemental Table 7). For each of 291 samples for which matching polyA and total RNA data was available, we calculated the log2 ratio cut-off that most accurately classified known polyadenylated (n=5,987) and non-polyadenylated (n=117) genes^22^ and subsequently applied it to establish the polyadenylation status of the stringent RNA Atlas genes (Supplemental Table 2, see Methods for details). Only genes with at least 10 counts were included in the analysis. As expected, most mRNAs (90%) were classified as polyadenylated (majority vote across all samples, Figure 3C). For lincRNAs and asRNAs, the fraction of polyadenylated genes was 48% and 63% respectively. Notably, up to 75% of the novel single-exon lincRNAs were classified as non-polyadenylated (Figure 3D), demonstrating the added value of total RNA-sequencing to detect this specific RNA biotype. To empirically validate our polyA-status assessment methodology, we established a polyA-minus RNA-sequencing protocol by depleting polyadenylated transcripts from total RNA libraries and applied it to two RNA Atlas cell lines. Non-polyadenylated genes showed significantly higher polyA-minus/polyA count ratios than polyadenylated genes (P < 1 x 10^−10^, Wilcoxon signed-rank test) supporting our proposed classifications (Figure 3E).

To study potential changes in gene polyadenylation status across samples, we focused our analysis on genes with at least 100 counts in 20 or more samples. Log2 count ratio distributions across samples suggested shifts from a polyadenylated to a non-polyadenylated state for a subset of genes in each biotype category (Figure 3F). Genes that were classified at least twice as non-polyadenylated and at least twice as polyadenylated were enriched among the lincRNA (59%) and asRNA (64%) biotypes compared to the mRNA biotype (43%). To evaluate what may be driving these shifts, we selected the more extreme cases based on log2 count ratios in each class (< −4 in at least 2 non-polyadenylated samples, > 0 in at least 2 polyadenylated samples). This selection included 83 mRNAs, 36 lincRNAs and 39 asRNAs (Figure 3G, Supplemental Table 8). We found that variable polyadenylation status was driven by differential expression of alternatively polyadenylated isoforms in 57 (36%) of these genes. One example is the histone gene HIST1H2BD that is transcribed into a single exon non-polyadenylated mRNA in the ALL-SIL cell line and a two-exon polyadenylated mRNA in prostate tissue (Figure 3H). Of note, the remaining 103 genes did not show any obvious changes in isoform usage, suggesting the existence of also other mechanisms causing differences in polyadenylation status (Figure S9).

### RNA biotype expression reflects sample ontology

After establishing and annotating a stringent transcriptome that covers the five primary RNA-biotypes, we analyzed RNA biotype expression data in relation to sample ontology and transcriptome biology. We first validated that the RNA Atlas expression data reflects a number of well-established features of the transcriptome such as non-coding RNA expression specificity, imprinting and cancer fusion gene expression. As expected, we observed a strong enrichment of mRNA fusion genes in cancer cell lines compared to non-malignant cell types and tissues (Figure S10) and detected 20 known imprinted genes that featured consistent mono-allelic expression over the large majority of samples (Figure S11, Supplemental Table 9). When evaluating expression-specificity for the various biotypes, the RNA Atlas expression data verified that non-coding RNAs were expressed in a more tissue-specific manner than mRNAs (Figure S12). Notably, when correcting for differences in expression abundance between RNA biotypes, lincRNAs and asRNAs remained more specific than mRNAs while circRNAs showed similar specificity (Figure S12). When selecting 1,320 tissue-specific RNAs from external datasets^23^, 96.14% were cross-validated in the RNA Atlas dataset, independently validating the RNA Atlas gene expression profiles (Figure S13). Based on these observations, we concluded that the RNA Atlas expression data indeed reflects the known characteristics of the transcriptome.

We then calculated expression-based distances between samples to evaluate associations between expression data and sample ontology for each of the biotypes. With this analysis, we evaluate if the large collection of novel genes that were identified (such as the single exon lincRNAs and novel miRNAs) are expressed in a non-random manner, and thus reflect underlying sample-ontology relationships. We first focused our analysis on 4 major cellular subtypes, i.e. epithelial cells (n=21), endothelial cells (n=25), fibroblasts (n=33) and mesenchymal cells (n=8). We reasoned that the transcriptional profiles of cells within each subtype should be closely related, and distinct from transcriptional profiles of the other cell types in the RNA Atlas dataset. Two-dimensional clustering of all cell types based on mRNA expression data confirmed that cell types within each subtype were indeed more closely associated (Figure 4A).

Consequently, mRNA expression-based distances between samples within a subtype (intra-distance) were significantly smaller than expression-based distances between these samples and all other samples in the dataset (inter-distance, P < 0.05, paired T-test, Figure 4B). This was observed for all 4 subtypes and was most pronounced for endothelial cells for which the median inter-distance was 2.3-fold higher than the median intra-distance. When calculating intra- and inter-distances for the other RNA biotypes, lincRNAs, asRNAs and miRNAs all showed inter-distances that were significantly higher than intra-distances (Figure 4C). More importantly, when calculating these distances using the novel miRNAs or single-exon lincRNAs, we observed similar patterns, suggesting these genes are also expressed in a coordinated manner (Figure 4D-E). This is supported by the fact that a subset of novel miRNAs (n=68) and single exon lincRNAs (n=197) are significantly upregulated (log2 fold change of at least 3 and a Benjamini-Hochberg based FDR lower than 0.01) in individual cell subtypes and demonstrate subtype-specific expression profiles (Figure 4F-I). Surprisingly, for circRNAs, we did not observe a substantial difference between inter- and intra-distances (Figure 4C). To further validate these findings, we repeated our analysis, focusing on the cancer cell lines within each cancer type. Inter-distances (expression distances between cell lines from the same cancer-type and cell lines from other cancer-types) were significantly higher than intra-distances (expression distances among cell lines from the same cancer-type) for mRNAs, asRNAs, lincRNAs and miRNAs but not for circRNA (Figure S14). Significant differences were also observed for the novel miRNAs and single-exon lincRNAs (Figure S14), confirming our observations on the biological subtypes. Moreover, we identified several novel miRNAs (n=154) and single-exon lincRNAs (n=1,006) that were significantly overexpressed in individual cancer types (Figure S14).

These results indicate that the genes in the RNA Atlas transcriptome, and more specifically the novel miRNAs and single-exon lincRNAs, show expression patterns that closely reflect sample ontology relationships. Furthermore, these non-random expression patterns again support that the single-exon lincRNAs do not derive from DNA fragments contaminating the RNA-sequencing libraries.

Both for the cell subtypes and the cancer types, circRNA based expression distances did not differ between related and unrelated samples (Figure 4C and Figure S14C). Because of technical limitations (circRNAs can only be quantified using reads spanning the back-spliced junction), the number of available reads that quantify circRNA expression is typically low, and thus inherently more variable compared to linear RNAs. When focusing our analysis on more abundant circRNAs, we indeed observed an increase in the difference between the inter and intra-distance (Figure S15). That increase was proportional to the abundance of the selected circRNAs and only the 1% most abundant circRNAs produced results similar to those observed for the linear RNAs. Overall, 34 and 108 circRNAs were significantly upregulated (log2 fold change of at least 3 and a Benjamini-Hochberg based FDR lower than 0.01) in one of the 4 major cell subtypes or 10 cancer types, respectively (Figure S15).

### Total RNA transcriptomes facilitate the use of intron expression profiles to study regulatory modalities

Our total RNA sequencing profiles provide broad coverage of expressed pre-mRNA introns (Figure 5A). Consequently, we were able to estimate both intronic and exonic expression for most widely expressed transcripts, and we used these profiles as surrogates for pre-mRNA and mRNA expression profiles, respectively (Supplemental Tables 10-12). Analysis of these pre-mRNA and mRNA expression profile estimates suggested a universally significant but less than a perfect concordance for most transcripts (Figure 5B). Interestingly, pre-mRNA and mRNA expression deviated significantly more for genes with longer 3’ UTRs (P < 8 x 10^−37^), which can be explained by tighter 3’UTR-mediated post-transcriptional regulation^24^. Namely, transcriptional regulation is expected to affect both pre-mRNA and mRNA expression profiles while post-transcriptional regulation should be evidenced by deviations between these profiles. To further study the effects of transcriptional and post-transcriptional regulation, we collected verified transcription factor (TF) and miRNA targets for 210 TFs and 226 miRNAs (Supplemental Table 13). TF expression had significantly higher correlations with both the target pre-mRNA and mRNA expression than with the target mRNA/pre-mRNA-ratio (further referred to as the m/p-ratio). In contrast, and as expected, miRNA expression had significantly higher correlations with the target m/p-ratio than with pre-mRNA and mRNA expression of the target (Figure 5C-D). These observations verify that mRNA and pre-mRNA estimates can be exploited to study transcriptional and post-transcriptional regulation. Moreover, they provide a means to evaluate the miRNA-like behavior of the novel miRNAs identified in this study in relation to their predicted targets. However, extending this observation to predicted TF and miRNA targets required accurate and dataset-specific regulator-target predictions (Supplemental Table 14) because sequence-based target predictions alone showed little evidence of differences between pre-mRNA and m/p-ratio correlations (Figures 5E and S16). Indeed, analyses focusing on miRNAs with predicted targets by LongHorn^25^ suggested that m/p-ratio correlations for miRBase miRNAs were significantly higher than pre-mRNA correlations. Note that LongHorn uses mRNA expression estimates to predict interactions, and, consequently, LongHorn-predicted regulator and target mRNA profiles—but not m/p-ratios—are expected to be correlated. Similar to miRBase miRNAs, novel RNA Atlas miRNAs had significantly higher m/p-ratio than pre-mRNA correlations (Figure 5F, P < 6 x 10^−8^, Kolmogorov-Smirnov test).

We set out restrictive criteria to identify individual miRNAs with significantly better m/p-ratio than mRNA and pre-mRNA profiles. Namely, only miRNAs with at least one predicted target with adequate pre-mRNA expression profiles in all the RNA Atlas samples were included in the analysis. In total, 719 miRBase miRNAs and 512 novel RNA Atlas miRNAs satisfied this requirement. Of these, 672 miRBase miRNAs (93%) and 436 novel RNA Atlas miRNAs (85%) had at least one target for which we observed a higher correlation between its expression profile and its target’s m/p-ratio profiles than its target’s mRNA and pre-mRNA profiles. However, to test that the expression profiles of miRNAs are significantly more likely to have stronger correlation with their target’s m/p-ratios, we required that a significantly greater number of predicted targets have higher miRNA to m/p-ratio correlations (P < 0.05 by t test). In total, of the 699 miRBase miRNAs and 485 RNA Atlas miRNAs that had multiple interactions with target pre-mRNA expression available, 198 miRBase miRNAs (28%) and 99 RNA Atlas miRNAs (20%) satisfied this requirement (Supplemental Table 4). Our analysis suggests that these 198 miRBase miRNAs and 99 RNA Atlas miRNAs are functionally regulating multiple post-transcriptional targets across multiple samples in our dataset. The identity of these miRNAs and their targets in given in Supplemental Table 15.

### Evidence for transcriptional and post-transcriptional regulation by long noncoding RNAs

Our ability to distinguish transcriptional from post-transcriptional regulation could also provide insight into lncRNA function. We first applied LongHorn^25^ to infer regulatory networks downstream of RNA Atlas lncRNAs, including single- and multi-exon lincRNAs, asRNAs and circRNAs. LongHorn (Figure 6A) predicts lncRNA targets by evaluating them as modulators of either transcriptional regulation - where lncRNAs are modeled to alter TF regulation as co-factors, guides, or decoys - or of post-transcriptional regulation as miRNA and RNA-binding protein decoys^26–29^ We then evaluated whether LongHorn predictions were supported by the expected correlations between regulator expression (i.e. TF or miRNA) and target mRNA, pre-mRNA, and m/p-ratio estimates. For each LongHorn prediction, we selected TF-target interactions based on ENCODE ChIP-seq^30^ and TF binding motifs and miRNA-target interactions based on sequence analysis^25^. Analogously to verified TF-target interactions (Figure 5C) and for all types of lncRNA modulators considered, pre-mRNA and mRNA correlations were consistently higher than m/p-ratio correlations for predicted TF-target interactions (Figure 6B). Note that predicted TF-target interactions with no evidence of lincRNA modulations did not have substantial differences between pre-mRNA and m/p-ratio correlations (Figure S16). Similarly, all types of lncRNA mediated miRNA-target interactions showed higher m/p-ratio correlations than mRNA or pre-mRNA correlations (Figure 6C); see Supplemental Tables 16-17 for the complete data.

**Figure 6.**
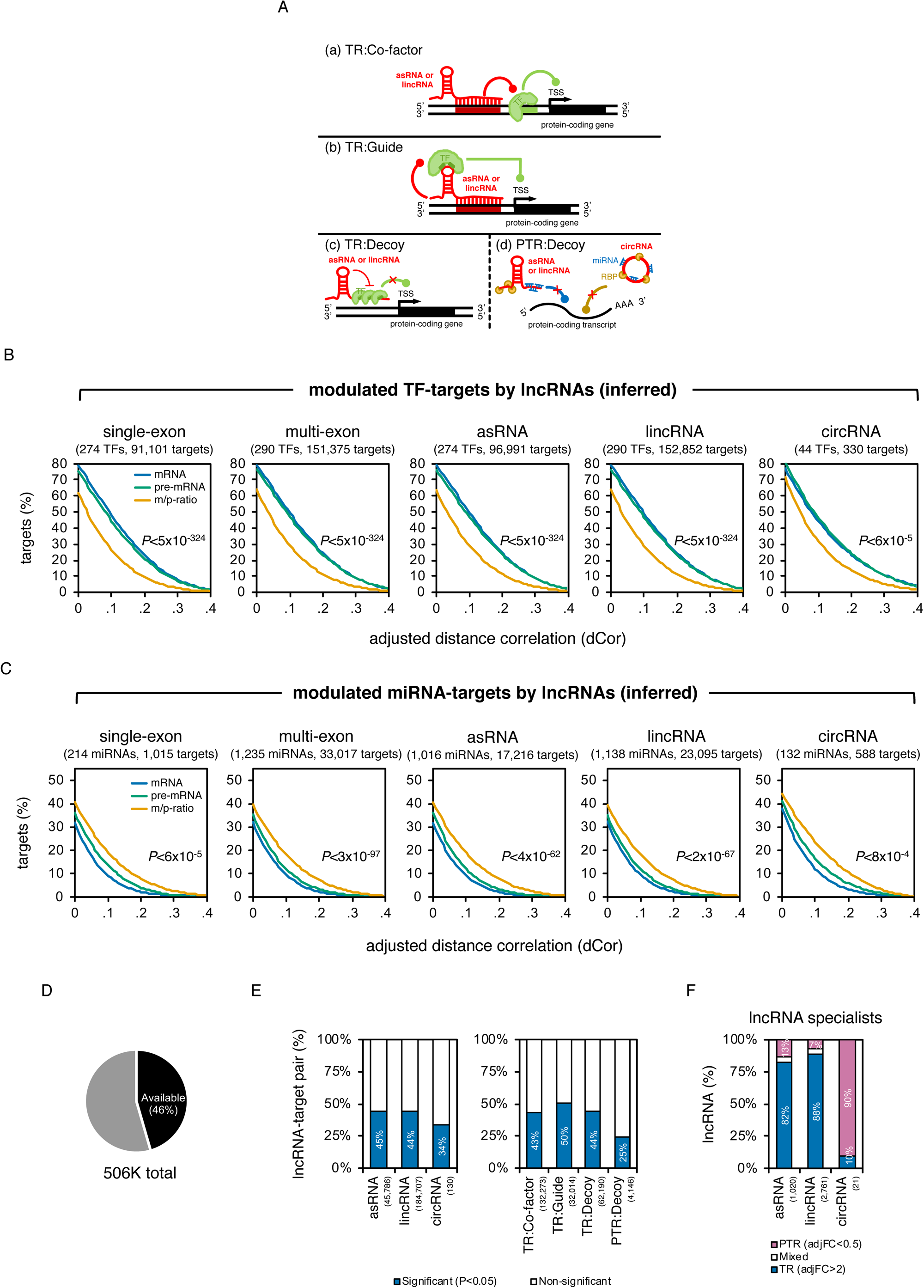
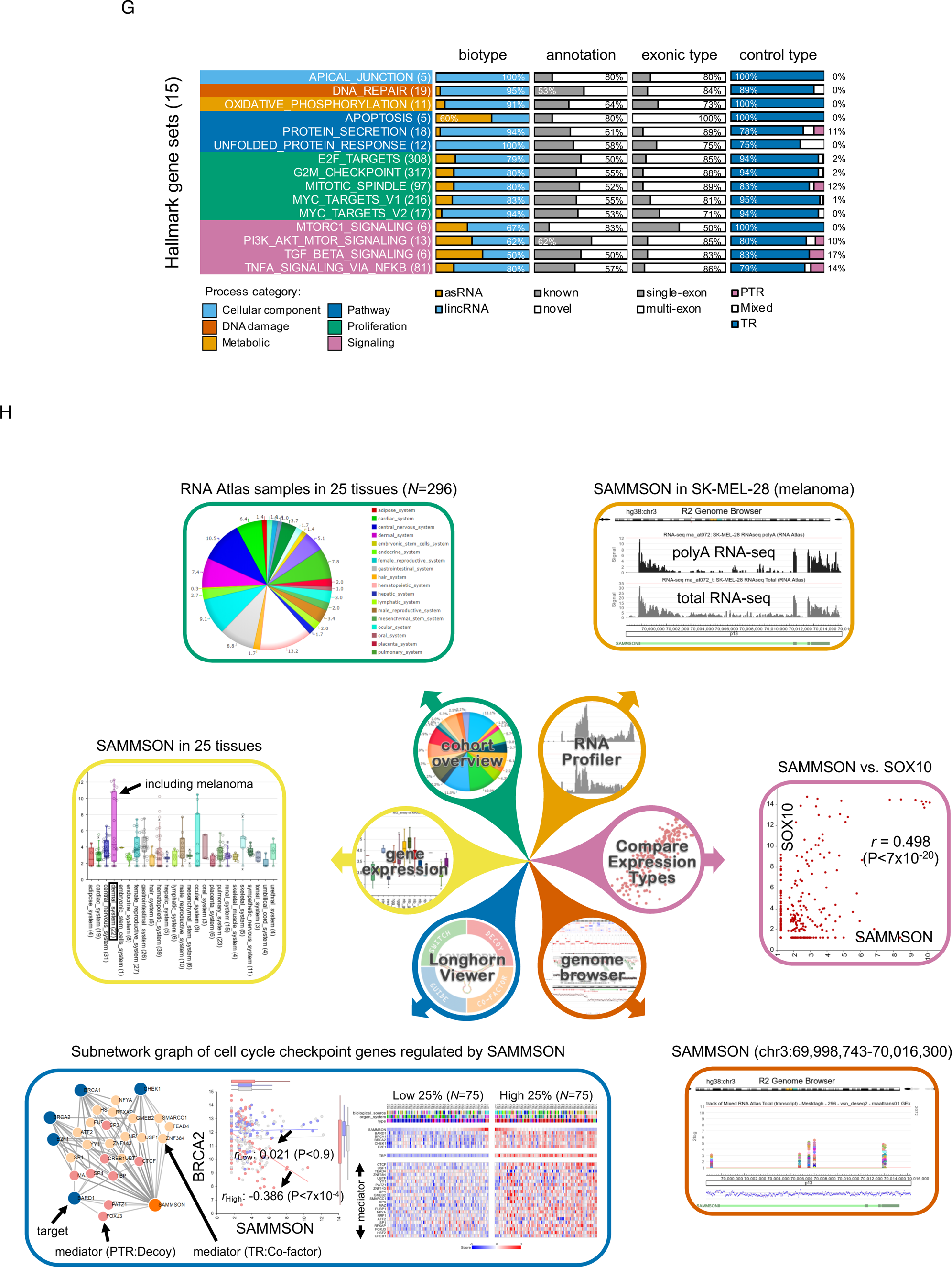
Evidence for regulation by lncRNAs. (A) LongHorn inferred interactions by evaluating four distinct models for lncRNA regulation, including transcriptional regulation (TR) by lncRNAs as co-factors, guides, or TF decoys, and post-transcriptional regulation (PTR) as decoys for miRNAs and RNA-binding proteins. (B) Predicted TF-target interactions with no supporting evidence from expression did not show correlation differences between TF-target pre-mRNA and m/p-ratio, see Figure S16. However, evidence for lncRNA regulation, including regulation by single- and multi-exon lncRNAs, antisense, intergenic, and circular lncRNAs, resulted in TF-target pre-mRNA profiles that were significantly better correlated than TF-target m/p-ratio profiles; the number of tested TFs and interactions for each lncRNA biotype is given in parentheses. Similarly, (C) miRNA-target interactions modulated by single- and multi-exon lncRNAs, antisense, intergenic, and circular lncRNAs showed higher miRNA-target m/p-ratio correlations. (D) LongHorn predicted 506K lncRNA-target interactions, and among these, 46% had sufficient intronic and exonic read counts to evaluate m/p-ratio in all profiled samples. (E) Proportion of interactions for which regulator-target m/p-ratio correlations were significantly different from regulator-target pre-mRNA correlations. Results are shown by the biotype of modulating lncRNAs (left) and the type of regulation for these interactions (right). The regulators were either TF or miRNAs mediating the lncRNA-target interactions. p-values were estimated using the two-sample student’s *t*-test. (F) LongHorn identified lncRNAs that are enriched for predicted transcriptional or post-transcriptional interactions, or for both, relative to other lncRNA species. asRNAs and lincRNAs were more likely to be identified as transcriptional regulators, while circRNAs were more likely to be identified as post-transcriptional regulators; each of these lncRNA was required to have at least ten LongHorn-inferred targets; the number of lncRNAs included in each category is given in parenthesis. (G) Fifteen MsigDB’s Hallmark Gene Sets were predicted to be significantly regulated by at least five lncRNAs (FDR-adjusted Fischer Exact Test FDR-FET<0.01). In particular, lncRNAs, the majority of which are RNA Atlas predicted, were predicted to target proliferation and signaling pathways; the total number of regulating lncRNAs in each pathway is provided in parentheses. (H) The full lncRNA-target prediction data is available for download and analysis on the R2 platform. R2 also allows the analysis and visualization of lncRNA abundance and regulatory network modules predicted by LongHorn. We show an example of an R2 analysis for the lncRNA SAMMSON.

These observations thus suggest that all lncRNA biotypes, including single-exon lincRNAs, are effectively altering TF and miRNA regulation. In addition, for many lncRNA species, the analysis pointed to lncRNA specialization as either transcriptional or post-transcriptional regulators. In total, we predicted 506,659 lncRNA-mediated interactions and targets included in 230,623 of these (Figure 6D) had pre-mRNA, mRNA, and m/p-ratio expression estimates in all samples. Nearly half of these interactions showed significant differences between pre-mRNA and m/p-ratio correlations. However, there were fewer significant differences for interactions that were modulated by circRNAs and for post-transcriptional decoys (Figure 6E). Indeed, an analysis of lncRNA regulatory models inferred by LongHorn suggested that circRNAs are predominantly post-transcriptional decoys, while other lncRNAs predominantly modulate transcription (Figure 6F). These results are in line with earlier observations demonstrating the enrichment of circRNAs in the cytoplasm and lincRNAs in the nucleus^31–33^. In total, 981 single-exon lincRNAs (75% of lncRNAs that modulated at least one interaction for which a m/p-ratio was computed) had significant differences between pre-mRNA and m/p-ratio. An additional 2,305 multi-exon lincRNAs showed similar behavior.

Nearly 4000 lncRNAs were associated with ten or more LongHorn-inferred targets, and the majority of these lncRNAs were never previously cataloged. Our analysis suggests that many of these lncRNAs preferentially target key pathways in disease and development (FDR-adjusted Fisher Exact Test FDR-FET<0.01). To analyze this further, we cataloged lncRNAs according to their predicted targets’ enrichment in hallmark pathways^34^ and their specialization as transcriptional or post-transcriptional modulators. In total, 15 pathways were enriched in targets from at least five lncRNAs (Figure 6G). Our analysis suggests that, overall, lncRNAs preferentially target proliferation and signaling pathways. The full analysis is given in Supplemental Table 18 and pathway enrichments for twenty-five known and novel lncRNAs are depicted in Figure S17. The full lncRNA-target prediction data is available for download and analysis on the R2 platform (r2.amc.nl). R2 allows for comparing profiles across technologies, visualizing lncRNA expression across samples, and studying regulatory network modules predicted by LongHorn, including analyzing correlations between lncRNAs, TFs, miRNAs, and predicted targets. Figure 6H depicts an example of the R2 analysis modules using the lncRNA SAMMSON^15^.

## Discussion

By applying three complementary RNA-sequencing technologies on a heterogeneous collection of tissues, cell types and cell lines, we assembled a very comprehensive human transcriptome covering all major RNA biotypes. While our effort complements other consortium-based efforts aimed at generating human expression atlases^4–12^, it also vastly extends beyond their scope in various ways. This is mainly achieved through the integration of a total RNA-sequencing methodology, providing a layer that is missing in the hitherto available compendia, on top of polyA enriched sequencing. Not only did the total RNA sequencing component of the RNA Atlas dataset reveal novel non-polyadenlylated lincRNAs and circRNAs, it also enabled us to determine alternative transcript polyadenylation status and quantify intronic RNA abundance levels. The latter is crucial to distinguish transcriptional from post-transcriptional regulation, a concept we have exploited in various ways. First of all, we have used it to reduce the more than 3000 newly assembled miRNAs into a stringent set of 99 miRNAs with strong evidence of post-transcriptional regulation of gene expression in multiple samples in our dataset. While we imposed minimal abundance criteria for the novel miRNAs, we did not implement other criteria defined in the miRBase high-confident checklist^35^ as opposed to a similar effort published earlier by the FANTOM consortium^8^. Notably, several miRBase miRNAs have validated target genes but do not adhere to the high-confident criteria, which are mainly related to sequence characteristics. We therefore reasoned that miRNA-like behavior (i.e. post-transcriptional regulation of target gene expression) outweighs sequence characteristics when defining stringent miRNAs. Second, we have integrated total RNA-sequencing intron and exon expression levels with the LongHorn algorithm^25^ to infer non-coding RNA regulatory interactions.

The integration of miRNA and whole-transcriptome profiling across a variety of tissues enabled unique analyses that, in turn, allowed us to evaluate RNA Atlas predicted ncRNA species for functional evidence. Consequently, in addition to identifying new miRNAs and lncRNAs, we were also able to collect multiple lines of evidence for their functional relevance in human cells and to interpret their function through both transcriptional and post-transcriptional regulatory interactions. The resulting RNA Atlas dataset and analysis products serve as a resource to mine the expression and regulatory landscapes of multiple RNA biotypes and contains a unique collection of non-coding RNAs together with their functional interpretation. Dedicated experimental validation studies based on genetic perturbations coupled to phenotypic or molecular readouts should follow to evaluate ncRNA function^36^ in each studied context. Moreover, we envision that the non-coding RNA regulatory interactions that are presented will serve as a starting point for follow up studies to gain insights into the mode-of-action of hundreds of ncRNAs.

## Methods

### Sample cohort

A total of 300 human samples were used in this study, including 45 tissues, 162 cell types and 93 cell lines, of which 89 are cancer cell lines derived from 13 different types of cancer (Supplemental Table 1). RNA of individual cell types was obtained from ScienCell Research Laboratories or isolated from cell types collected at Ghent University Hospital. RNA from collected cell types and (cancer) cell lines was isolated using the miRNeasy kit (Qiagen) according to the manufacturer’s instructions. RNA samples from normal human tissues were obtained from Ambion and Biochain.

### Library prep and sequencing

For each RNA sample, three different strand-specific RNA libraries were prepared. Small RNA libraries were generated using the TruSeq Small RNA library prep kit (Illumina) according to the manufacturer’s instructions, using 750 ng input RNA. Library size selection was performed using a Pippin Prep device (Sage Science). Total RNA libraries were generated using the TruSeq stranded Total RNA library prep kit with Ribo-Zero Gold (Illumina) according to the manufacturer’s instructions using 1 µg of input RNA. PolyA RNA libraries were generated using the TruSeq Stranded mRNA library prep kit (Illumina) according to the manufacturer’s instructions using 1 µg of input RNA. Small RNA libraries were pooled (volume-based pooling, 48 libraries per pool) and pools were quantified using the High Sensitivity dsDNA assay on a Bioanalyzer device (Agilent). Poly A and total RNA library pools were quantified using the Standard Sensitivity NGS Fragment Kit (Catalog #DNF-473) on a Fragment Analyzer (Advanced Analytical, Ankeny IA, USA). Small RNA library pools were sequenced on a NextSeq500 instrument (Illumina) using a high output flow cell, 76 cycles. Pooled PolyA and Total RNA libraries were sequenced on a HiSeq 4000 instrument (Illumina) with paired-end 76 cycle reads.

### Transcriptome assembly from polyA and total RNA-sequencing libraries

PolyA and total RNA reads were mapped to the hg38 reference genome (primary assembly, canonical chromosomes, repeats from RepeatMasker and Tandem Repeats Finder soft masked http://hgdownload.soe.ucsc.edu/goldenPath/hg38/bigZips/) with TopHat^37^ v2.1.0 (Bowtie2^38^ v2.2.6) using the --no-coverage-search option and --library-type=fr-firststrand. The Ensembl^17^ transcriptome (v86) was provided to guide the mapping of reads to known transcripts first. All other parameters were set to default values. Transcriptomes were assembled in each sample and each library type separately using StringTie^39^ (v1.3.3). Default parameters were used and the Ensembl^17^ reference annotation (v86) was supplied to guide the assembly.

All individual transcriptomes were merged together with the reference Ensembl transcriptome (v86) using StringTie merge applying a cutoff of 1 TPM and minimum transcript length of 200 nucleotides, with all other parameters set to default values.

Cuffcompare^40^ (v2.2.1) was used to compare the newly assembled transcripts with the reference annotation. Novel transcripts with classification codes other than ‘x’, ‘u’,’j’ or ‘=’ were removed, (this included 20,539 transcripts with codes ‘c’, ‘e’, ‘I’, ‘o’, ‘p’ or ‘s’) as well as transcripts spanning 2 or more known, non-overlapping, adjacent loci.

We then calculated the overlap of all known and novel exons with repetitive elements in the genome using BEDTools^41^ (intersect). The repeats regions were retrieved from the UCSC Table Browser^42^ (Group: Repeats; Track: RepeatMasker). The fraction of exonic sequence overlapping repeats was computed for each gene. Novel non-coding single-exon genes with less than 200 consecutive non-repeat nucleotides were filtered out. Regions overlapping repeats within the remaining non-coding single-exon genes were hard-masked (bases were converted to Ns) using BEDTools^41^ (v2.27.1, maskfasta). After these filtering steps, the polyA/total RNA-sequencing derived transcriptome contained 422,083 transcripts including all transcripts in Ensembl v86 annotation. This transcriptome was quantified using Kallisto quant^43^ (flag --rf-stranded and all other parameters set to default values) across all polyA and total RNA-sequencing libraries.

After quantification, novel genes and Ensembl genes belonging to the biotypes ‘protein_coding’, ‘antisense’ and ‘lincRNA’ were retained. Because of previous filtering steps at transcript level, novel genes are either intergenic (lincRNA) or antisense (asRNA) to reference genes. Known and novel genes with expression levels below 0.1 TPM in all samples were removed.

### Selection of the stringent RNA Atlas transcriptome

Independent evidence for transcription of the mRNA, lincRNA and asRNA genes in the RNA Atlas transcriptome was obtained by integrating results from cap analysis of gene expression (CAGE) sequencing from the FANTOM consortium^6^ and various chromatin states from the Roadmap Epigenomics Project^18^. The following chromatin states, indicative for active transcription, were considered: active transcription start site (1_TssA), transcription (5_Tx5, 6_Tx and 7_Tx3), transcribed and regulatory (9_TxReg), transcribed and enhancer (10_TxEnh5 and 11_TxEnh3), active_enhancer (13_EnhA1 and 14_EnhA2) and bivalent_promoter (23_PromBiv)^18^. For each TSS of genes with expression values greater or equal to 0.1 TPM in at least one tissue from the RNA Atlas and not being part of chromosome Y (chromatin states did not include information for that chromosome), we used the Zipper plot approach^44^ to retrieve the closest CAGE-seq^6^ and chromatin state^18^ peak across all samples from the FANTOM5 and Roadmap Epigenomics project, respectively. We defined the stringent gene set based on presence of the aforementioned peaks within 500 nucleotides upstream or downstream the TSS and further classified the genes across the different RNA biotypes into four categories: 1) evidence at both DNA (chromatin state) and RNA (CAGE-seq) level, 2) evidence at RNA level only, 3) evidence at DNA level only, 4) no evidence. Only genes belonging to one of the first 3 categories were retained in the stringent RNA Atlas transcriptome.

We also included 754 genes in the stringent set that do not present close association to any CAGE-peak or chromatin state peaks but whose expression levels are higher than the median expression levels for genes that present both levels of evidence. In more detail, we retained genes with no independent evidence if their mean expression across samples in polyA or total RNA was higher than the median value of the mean expression across samples for genes with both levels of evidence (4.67 TPM and 2.20 TPM, for polyA and total RNA data, respectively), or if their maximum expression across samples in polyA or total RNA was higher than the median value of the maximum expression across samples for genes with both levels of evidence (58.63 TPM and 30.23 TPM, for polyA and total RNA data, respectively). This set includes 510 mRNAs (508 known and 2 novel), 27 asRNAs (17 known and 10 novel) and 208 lincRNAs (95 known and 113 novel).

### Evaluation of coding potential

To assess the protein-coding potential of the new transcripts, two algorithms were used: The Coding-Potential Assessment Tool^45^ (CPAT, version 2.0.0) and PhyloCSF^46^ (obtained from https://github.com/mlin/PhyloCSF January 18, 2015). The CPAT code was slightly modified so that the predicted ORF sequence is returned in the output. CPAT was run using the provided hexamer table and logit model. The recommended probability cutoff of 0.364 was used to identify putative coding ORFs.

Since the PhyloCSF pipeline is based on the GRCh37/hg19 reference genome, all genomic coordinates were first converted using the liftOver tool on the UCSC Genome Browser website^42^. To run PhyloCSF, whole-genome alignments of 46 species are obtained from the UCSC Genome Browser website^42^ and processed using the Phylogenetic Analysis with Space/Time Models package^47^ (PHAST, v1.4) into the required input format. PhyloCSF was run in 3 reading frames using the ATGStop setting, all ORFs of at least 10 codons were considered. A cutoff score of 60.7876, based on benchmarking with Ensembl (v90)^48^ transcripts, was used to identify putative coding ORFs. In total, 242 novel genes (188 novel stringent) had at least one isoform scored as protein-coding by both tools. We labeled these genes as novel protein-coding. All other novel genes were annotated as lincRNA or antisense.

### Mass spectrometry validation

Mass spectrometry validation of the novel predicted proteins was conducted on the draft map of the human proteome dataset^49^. Briefly, this dataset consists of deep proteomic profiling of 17 adult tissues, 7 fetal tissues, and 6 purified primary hematopoietic cells. Raw files were obtained from the PRoteomics IDEntifications (PRIDE) database^50^ (project PXD000561) and converted to Mascot generic format (MGF) using the msconvert tool in the ProteoWizard package^51^.

Analysis of the tandem mass spectrometry data is performed using Ionbot (unpublished, based on the work of C Silva et al.^52^, https://ionbot.cloud), a sequence database search tool based on machine learning capable of performing rapid open modification and open mutation searches. Ionbot was used under a beta-tester version supplied by Sven Degroeve and Lennart Martens (Ghent University, VIB). Briefly, peptide databases are created as a full *in silico* trypsin digestion (allowing up to one missed cleavage) of the protein sequence dataset consisting of all human proteins in the Uniprot in the UniProtKB database^53^ (Swiss-Prot subset, 21,008 proteins) and the CPAT and PhyloCSF predicted proteins. Decoy peptides are obtained by applying the same digestion on the reversed target proteins. Ionbot was run in the open modification and open mutation mode. In addition, carbamidomethylation of cysteine was set as fixed and oxidation of methionine as variable modification. The false discovery rate (FDR) was estimated with the target-decoy approach. Only peptide-spectrum matches with an estimated FDR below 1% were retained.

### Identification and quantification of circRNAs

CircRNAs where identified from total RNA-sequencing data using two independent workflows, find_circ2^54^ (n=85,470) and CIRCexplorer2^55^ (n=204,857) using genome build hg19. For downstream analysis, the mean circRNA count across methods was used. Only circRNAs identified by both tools (n=62,832) and with mean counts between tools higher than 4 in at least one sample were retained (n=38,030). A circRNA was annotated as known if its back-splice position was present in the circBase database^19^. Genomic positions of 38,023 circRNAs were successfully converted to hg38 coordinates using the liftOver tool (UCSC)^42^. The back-splice acceptor and donor sites from each circRNA were annotated relative to other linear-splice sites and gene coordinates from mRNAs, asRNAs and lincRNAs. CircRNAs with a back-splice acceptor and donor site overlapping genes in the stringent RNA Atlas transcriptome were retained as stringent circRNAs (n=37,140).

### Flanking intron length analysis

We compared the length of the introns (both upstream and downstream) which flanked the circRNAs, to introns from genes that do not produce a circRNA isoform. The flanking introns were found to be unusually long when compared to the non-circRNA introns as shown in Figure S3). The median length for the flanking introns was 6304 bp compared to the median value for non-circRNA introns which was observed to be 1041 bp. Statistical significance of the difference was assessed with the Wilcoxon signed-rank test. Box plots were drawn in R to display the results.

### miRNA identification and quantification

Reads from small RNA-sequencing libraries were processed with the FASTX-Toolkit^56^ (v0.0.14) to remove adapters, filter reads by quality (a quality score of at least 20 in 80% of the sequence was required) and collapse non-unique reads. Processed reads were then mapped against the hg38 genome with Bowtie^57^ (v1.1.2) allowing no mismatches in the first 25 bases of the read (-n = 0 and l = 25) and using the “--best” option to force reporting of up to 10 (k = 10) best alignments (all other parameters were set to default values). Novel miRNAs were predicted with miRDeep2^58^ (v2.0.0.8), using mapped reads per sample and all human miRNAs in miRBase^35^ v22 as input. Novel miRNA predictions with non-zero estimated probability were aggregated across samples, retaining only the prediction with maximum counts from both mature forms in a given sample in cases of predictions with partially overlapping coordinates. Reads mapped to the aggregated newly predicted miRNAs and human miRNAs from miRbase v22 were then quantified across all samples. For each miRNA, counts from the canonical mature form and non-canonical mature forms (isoMiRs) were aggregated. Only miRNAs with 10 or more counts in at least one sample were retained. Filtered miRNAs and their precursors were assigned unique names (RNAATLAS(p)MIR prefix followed by sequential numbers from 1 to 6740, which is the number of unique filtered sequences including both mature and precursor miRNAs). Known and predicted precursors were annotated based on their overlap with exonic and intronic regions of mRNAs, asRNAs and lincRNAs or to intergenic regions.

### Genomic analyses of single-exon lincRNAs

The distance from each unique RNA atlas exon to its closest up- or downstream exon was retrieved with BEDTools^41^ (v2.27.1, bedtools closest -io -D a -s). A two-sample Wilcoxon signed-ranked test was used to compare the distances between single-exon lincRNAs and multi-exon lincRNA exons. Sequence motifs at the exon-intron boundary of multi-exon genes and exon-intergenic boundary of novel single-exon genes were determined by calculating the frequency of each nucleotide at each position of the region starting 3 bases upstream and ending 3 bases downstream of the boundary. This was done for multi-exon mRNA exons, multi-exon lincRNA exons and single-exon lincRNA exons. The information content was computed for each position and the relative frequencies of each base in each position were represented as a sequence logo with the R package ggseqlogo^59^. For strandedness analyses, we selected unique exons with no overlap with any feature on the opposite strand. Only exons with 10 or more counts on the correct strand in at least 1 sample were considered. The strandedness for each selected exon was defined using the sample with maximum normalized expression on the correct strand, as the percentage of reads mapping to the exon on the correct strand relative to all reads mapping to the exon regardless of the strand.

### RT-qPCR validation of single-exon genes

We performed qPCR validation of single-exon genes using RNA from two RNA Atlas cell lines, SK-N-BE(2)-C and IMR-32. We designed specific forward and reverse primers for the amplification of a total of 110 genes, including 80 novel single-exon genes, of which 48 were ubiquitously expressed, 17 specifically expressed in SK-N-BE(2)-C and 15 specifically expressed in IMR-32, as well as 30 known single-exon genes included as a control. Primers were designed using primerXL^60^ (Supplemental Table 19). For each gene in each sample, two qPCR reactions were performed, one on cDNA and one on RNA (to assess amplification of contaminating DNA). cDNA was produced using the iScript Advanced kit (Bio-Rad) with a mix of random primers and oligo-dT primers on 2 µg of input RNA and a reaction volume of 20 µl. All qPCR reactions were performed in a total volume of 5 µl containing 2.5 µl of SsoAdvanced mastermix (Bio-Rad), 2 µl of forward and reverse primers (5 µM) and 0.5 µl of cDNA (10 ng/ul) or the equivalent mass of RNA. Reactions were run on a LightCycler 480 (Roche) in 384 well-plates using the following thermal cycling protocol: 2 minutes enzyme activation at 95.0°C (temperature ramp rate of 4.8°C/s), followed by 45 cycles of 5 s at 95°C (temperature ramp rate of 4.8°C/s) and 30 s at 60°C (temperature ramp rate of 2.5°C/s). For melting curve analysis, denature was performed at 95°C for 5 s (temperature ramp rate of 4.8°C/s), followed by cooling to 60°C for 1 min (temperature ramp rate of 2,5°C/s), and then heating to 95°C at a ramp rate 0.11°C/s with 5 acquisitions/°C. Final cooling was performed during 3 minutes at 37.0°C (temperature ramp rate of 2.5°C/s).

### Analysis of polyadenylation status

For these analyses we used read count expression data obtained with HTSeq-count^61^ (v 0.11.0) from TopHat bam files. 291 samples for which we have expression data from both polyA and total RNA-sequencing libraries were used. Samples “RNAAtlas249” and “RNAAtlas251” were not included in these analyses because they had a very high fraction of mitochondrial reads in polyA RNA-sequencing libraries. First, we generated a list of known polyadenylated and non-polyadenylated genes based on Yang et al. (2011)^22^ by selecting those genes that were annotated as either polyadenylated or non-polyadenylated in both cell lines used in that study. To normalize counts between matching polyA and total RNA-sequencing libraries for differences in library size and library complexity, we calculated the mean count of the 900 most abundant known polyadenylated mRNAs in both libraries, and used the mean count ratio between libraries (polyA/total RNA) as a scaling factor. For most samples this ratio was below 1 and was thus used to scale counts in the total RNA library. In cases where this ratio was higher than 1, we used the inverse of this ratio to scale the polyA library. Scaling was done by subsampling counts from the relevant library to obtain similar counts for polyadenylated genes in both libraries. Polyadenylation status was determined as follows: Only genes with at least 10 counts in total RNA libraries were classified, otherwise their polyadenylation status was considered as ‘undetermined’ (n=3,071). Genes with 0 counts in polyA and at least 10 counts in total RNA were classified as non-polyadenylated. For genes with non-zero counts in polyA and at least 10 counts in total RNA libraries a classification approach was taken, based on the log2 ratio of counts between polyA and total RNA libraries. First, a sample-specific log2 ratio cutoff was determined based on the distributions of known polyadenylated and known non-polyadenylated genes^22^. From these, only polyadenylated genes with at least 10 counts in the polyA library and only non-polyadenylated genes with at least 10 counts in the total RNA library were retained. A sample-specific log2 ratio cutoff was determined by taking the value that maximizes the accuracy (number of true polyadenylated genes + number of true non-polyadenylated genes)/(total number of genes)) of the classification of known genes into non-polyadenylated (log2 ratio below the cutoff) and polyadenylated (log2 ratio above the cutoff). Because the set of known polyadenylated genes is much larger than the set of known non-polyadenylated genes, we subsampled the polyadenylated genes to match the number of non-polyadenylated genes in order to obtain a balanced dataset. We repeated this approach 100 times, and took the mean selected cutoff across iterations. We then derived a general classification for each gene in the stringent set, by taking the majority vote across samples.

Heatmaps of gene polyadenylation across samples were plotted per biotype (mRNA, lincRNA and antisense). For this, the samples were first sorted based on a normalized log2 ratio obtained by subtracting the sample specific cutoff from the log2 ratios (to make them comparable across samples). Sorted samples were then binned in a total of 20 bins, and the mean corrected ratio for each bin was calculated and plotted in the heatmap.

To select genes with changing polyadenylation status across samples, we considered genes that are expressed in at least 2 samples with a corrected log2 ratio below −4 and a read count of at least 100 in total RNA, and expressed in at least 2 samples with a corrected log2 ratio above 0 and read counts of at least 100 in both total RNA and polyA. This resulted in 160 genes. To get insights into the factors driving the observed changes in polyadenylation status at gene level, we analyzed changes in expression levels of individual transcripts from these genes. Specifically, we retrieved the dominant transcripts in each library from the most extreme polyadenylated sample (i.e with highest log2 normalized ratio) and the most extreme non-polyadenylated sample (i.e with lowest log2 normalized ratio). We computed the fraction of total gene expression represented by the dominant transcripts and evaluated differences in dominance and fraction of expression between the polyadenylated and non-polyadenylated samples (Supplemental Table 8). By analyzing these parameters, we defined two cases in which the variability in gene-level polyadenylation across samples can be explained by differential expression of alternatively polyadenylated isoforms: (a)those genes that present a different dominant transcript in total RNA datasets from the polyadenylated and the non-polyadenylated samples, while presenting the same dominant transcript in the polyA and the total RNA datasets from the polyadenylated sample (n=48) or (b)those genes that present the same dominant transcript in total RNA datasets from the polyadenylated and the non-polyadenylated samples but whose fraction of total gene expression is lower in the polyadenylated sample compared to the non-polyadenylated sample, besides, the dominant transcript for the polyadenylated sample in its polyA dataset is not the same as the dominant transcript in its total RNA dataset (n=9).

### PolyA-minus sequencing

PolyA-minus libraries were generated for two RNA Atlas cell lines, SK-N-BE(2)-C and IMR-32 in duplicates. In brief, 500 ng of RNA was first depleted for rRNA using the Ribo-Zero Gold approach (Illumina) followed by the polyA selection procedure as implemented in the TruSeq mRNA library prep protocol (Illumina). Rather than discarding the polyA-minus fraction, 2 additional rounds of polyA selection were performed on that fraction, each time maintaining the polyA-minus fraction as input for the polyA selection step. The final polyA-minus fraction was concentrated using RNA Clean XP beads (Agencourt) before proceeding with library prep (according to the TruSeq mRNA library prep manual). In parallel, the polyA-plus fraction, obtained after the first polyA selection step, was also processed for library prep and sequencing. Libraries were quantified using qPCR (Kappa) and equimolarly pooled for sequencing on a NextSeq 500, high output flow cell, paired-end sequencing, 75 cycles per read (Illumina). Sequencing reads were mapped to the hg38 reference genome (primary assembly, canonical chromosomes, repeats from RepeatMasker and Tandem Repeats Finder soft masked http://hgdownload.soe.ucsc.edu/goldenPath/hg38/bigZips/) with TopHat^37^ v2.1.0 (Bowtie2^38^ v2.2.6) using the --no-coverage-search option and --library-type=fr-firststrand. The Ensembl transcriptome (v86)^17^ was provided to guide the mapping of reads to known transcripts first. All other parameters were set to default values. Read count expression data was then derived from the mapped reads using HTSeq-count^61^ (v0.11.0). Genes with at least 10 mean counts between replicates in either polyA-minus or polyA-plus libraries were selected and the ratio of polyA-minus versus polyA-plus counts scaled by library size was calculated. This ratio was then compared between genes that were previously classified as polyadenylated or non-polyadenylated in the corresponding sample based on the ratio of counts from polyA and total RNA libraries.

### Expression specificity

Expression specificity was computed for each RNA biotype and each sample type (cell types, cell lines and tissues) separately, using a specificity score based on the Jensen-Shannon divergence metric^62^. For mRNAs, asRNAs and lincRNAs, specificity was calculated using TPM values, for miRNAs and circRNAs, we used RPM values (reads per million).

These expression metrics are not directly comparable, because they come from library preparations that capture different sets of RNAs or, in the case of circRNAs, from reads mapping to one particular location in the transcript (i.e. the back-spliced junction) rather than the entire transcript (as is the case for mRNAs, lincRNAs and asRNAs). In order to directly compare expression levels between these biotypes, we quantified mRNA, lincRNA and asRNA expression based on junction reads. We compared the specificity score distributions of back-spliced junctions (circRNAs) and linear junctions (mRNAs, lincRNAs, asRNAs) before and after correcting for differences in expression abundance between these biotypes. For this, we subsampled subsets of splice junctions from each biotype so that the subsampled distributions of maximum expression values matched a common normal-like distribution with mean equal to the median of the mean values for the different biotype distributions and standard deviation equal to the mean minus one third of the smallest absolute extreme value (i.e. minimum and maximum values of the distributions) among all biotype distributions.

### Validation of tissue-specific RNAs from external datasets

Expression data from 23 tissues with matched RNA atlas tissues was retrieved from the Human Protein Atlas (HPA)^23^. 1,320 tissue-specific genes with an expression value of at least 5 TPM and a fold change of at least 10 between the first and the second tissue with highest expression values were selected within the HPA dataset. The selected HPA markers were considered as cross-validated in RNA atlas if they presented the highest expression in the same tissue. For all selected biomarkers the log2 fold-change between the expression in the matching tissue and the highest expression among the remaining 22 tissues was calculated.

### Fusion genes

Fusion genes were identified with FusionCatcher^63^ across all polyA-seq samples. In each sample, fusions labelled as probable false positives and fusions known to occur in healthy samples (Supplemental Table 20, codes 0 and 1) were filtered out. Also, the fusion transcripts were required to have zero ‘counts of common mapping reads’, i.e. reads that map on both partners, and a minimum of 4 unique reads mapping on the fusion junction. Finally, within each sample, transcript fusions were collapsed at gene level, i.e. if multiple junctions occurred at different joint points or reciprocally between the same pair of genes, they were counted only once, and the distribution of number of junctions per sample was compared between cell lines, cell types and tissues using two-sample Wilcoxon signed-ranked tests.

### Imprinting

To detect imprinting, data was first further processed according to^64^ which relies on Samtools (v0.1.19) for initial variant calling and genotyping (and sequencing error estimation) by SeqEM (v.1.0). Only variants present in dbSNP (version 150) were retained and insertions, deletions and loci corresponding with mutations from the Human Gene Mutation Database were removed. Detection of imprinting and other statistical analyses were performed in R (v3.3.2). Used filters were: coverage > 4, number of samples ≥ 75, minor allele frequency > 0.15, estimated sequencing error rate ≤ 0.035. As outlined earlier^64^, for the detection of imprinting across tissues per SNP, a mixture model of homozygous and heterozygous samples was fit to the RNA-seq data, with weights derived from Hardy-Weinberg equilibrium (HWE). The mixture model takes into account sequencing errors and partial imprinting. Unpublished before, also the degree of inbreeding in the underlying population is taken into account when estimating the fractions of heterozygous and homozygous loci, i.e. the weights of the mixture model. The degree of inbreeding is estimated as a hyperparameter, i.e. the median degree of inbreeding over all SNPs passing the quality filters (further described in^64^), leading to an estimate of 0.102. A likelihood-ratio-test is used to assess whether the model supports the absence of apparently heterozygous loci (which feature, on average, a 1:1 ratio of both alleles).

This methodology was applied on the Total RNA-seq samples from 203 tissue and cell type samples, excluding cancer/cell line samples given their frequent loss of imprinting^64^. Next to using Total RNA-seq, also the mRNA-seq dataset was queried (177 samples), whereas the coverage of the sRNA sequencing was too low to apply the methodology (data not shown). Other used filters were: goodness-of-fit > 1.2, symmetry > 0.05, median imprinting ≥ 0.8, estimated î ≥ 0.6, for more details, see^64^. Additionally, as we aimed to identify consistently imprinted loci, we focused on SNPs featuring a minimal difference between expected and observed heterozygous samples (based on SeqEM RNA-seq genotyping) of 30. A false discovery rate of 0.05 was used to call imprinting significant. We relied on RNA Atlas annotation, complemented by Ensembl annotation where relevant. In case of overlapping genes, the gene in which the SNP was located in an exonic region or UTR was selected. For additional validation, genotyping data from Illumina Human1M-Duo BeadChip for 10 cell lines (HEK-293T, SK-N-SH, A549, HL-60, K562, MCF-7, OVCAR-3, T-47D, JURKAT, H1 hESC), were downloaded from ENCODE for validation of imprinting in de cell lines. Note that these data had not been used during the screening phase. For virtually all sufficiently covered sample/gene combinations featuring heterozygous SNPs, at least one sample-SNP combination showed allelic expression patterns compatible with imprinting/mono-allelic expression.

### Expression-based distances and differential expression analysis

t-Distributed Stochastic Neighbor Embedding (t-SNE)^65^ was applied on the mRNA expression data for all cell types or cancer cell lines and the two first dimensions were used to plot a visual representation of the clustering. Weighted expression correlations (w_cor) for all pairs of samples were calculated for all RNA biotypes (using the cov.wt function in R^66^), using counts normalized by variance stabilizing transformation (VST, DESeq2)^67^ as input and the average of sigmoid transformations of VST normalized counts for both samples as weights. Expression distances (expr_dist) were derived from these values as: expr_dist = 1 - (w_cor + 1) / 2.

Expression distances were compared between cell types from 4 biological subtypes (epithelial cells (n=21), endothelial cells (n=25), fibroblasts (n=33) and mesenchymal cells (n=8)) or cancer cell lines from 12 cancer types (B-ALL (n=8), Breast cancer (n=6), Central Nervous System cancer (n=6), Colon cancer (n=7), Leukemia (n=6), Melanoma (n=9), Neuroblastoma (n=11), Non-Small Cell Lung cancer (n=9), Ovarian cancer (n=7), Prostate cancer (n=2), Renal cancer (n=8), T-ALL (n=8)) to measure inter- and intra-group distances. Paired t-tests were performed over the median intra and inter distances for each group.

Differential gene expression analyses (DESeq2)^67^ were performed to identify novel miRNAs, single-exon lincRNAs and circRNAs with a significant differential expression between cell subtypes or cancer types (Prostate Cancer and Leukemia were excluded from this analysis, for having only 2 cell lines belonging to the cancer type and for including a rather heterogeneous collection of cell lines, respectively). Those genes with a log2 fold change of at least 3 and a Benjamini-Hochberg based FDR lower than 0.01 where selected and expression data was visualized in heatmaps. For circRNAs, we repeated the previous analyses on cell types and cancer cell lines by using sequential subsets of top expressed circRNAs. For this, we sorted the circRNAs based on their mean back-spliced counts across samples within the sample sets used in each case and took the top 20, 10, 5, 3, 2, or 1 percent expressed circRNAs to calculate sample-sample distances and compared the results between subsets.

### Expression estimation of exons (mRNA), introns (pre-mRNA), and their ratios from total RNA-seq data

We sought to estimate the exonic (mRNA surrogate) and intronic (pre-mRNA surrogate) expressions of protein-coding transcripts. For each total RNA-seq BAM file profiled in RNA Atlas, the featureCounts v1.6.0 program^68^ was applied to enumerate read counts in each exon and intron regions.

Exon annotations were downloaded from the UCSC Genome Browser^42^ in December 2017 (track: NCBI RefSeq, table: refGene, assembly: GRCh38). We further extended exon boundaries by 10 base pairs to prevent exonic reads boundaries near the exon junctions from being considered as intronic reads^24^. After extension, regions that were within two consecutive exons of a protein-coding transcript but did not overlap with any exons of other coding- and non-coding transcripts were defined as intronic. We recorded exon and intron boundaries of each protein-coding transcript.

featureCounts was ran on exons and introns separately, with reads summarized at feature level, i.e., single exon or intron (argument: -f) and only primary alignments were counted (argument: --primary). Duplicate reads were excluded from the counting process (argument: --ignoreDup). Reads mapped to multiple genes (discordant reads) or locations (multi-mapping reads) were discarded.

Counts of reads matching entire exons and introns of the same transcript were used to represent its exonic (mRNA) and intronic (pre-mRNA) abundance, respectively. We required transcripts whose log2-transformed exonic and intronic expressions, after adding a pseudo-count of 8 to their raw read counts, are at least 5, i.e., 24 counts, in every RNA Atlas sample^24^. In total, 7,289 RefSeq transcripts—corresponding to 3,556 genes—were kept for analysis. To calculate exon/intron ratios (mRNA/pre-mRNA ratios or m/p-ratios in short), we used the following formula for each protein-coding transcript:

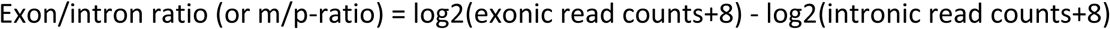

Each type of expression matrix, namely, mRNA, pre-mRNA, and m/p-ratio, was then separately normalized using quantile normalization over multiple RNA Atlas samples using quantilenorm routine in MATLAB; note that we used the median of the ranked values rather than the mean to perform normalization.

### Experimentally verified TF- and miRNA-targets

We compiled 6,535 experimentally verified TF-target pairs from three places including HTRIdb^69^ (version: 03/20/2014), TRANSFAC Professional^70^ (version: February 2013), and Table 3 from Whitfield et al.^71^ For pairs deposited at HTRIdb, we only included those verified by small and mid-scale techniques. To err on the conservative side and reduce false-positive predictions, we removed protein-DNA candidate interactions whose proteins are co-factors rather than TFs in TRANSFAC database. The list of 4,662 verified miRNA-targets with strong experimental evidence, such as western blot or reporter assay, were selected from miRecords^72^ (4/27/2013), TarBase^73^ v7, and TRANSFAC Professional ^70^(version: February 2013), miRTarBase^74^ v4.5, and Grosswendt et al., Table S2^75^. We can only keep a proportion of these interactions whose targets had sufficient exonic and intronic expression across all profiled RNA Atlas samples. In total, 2,349 and 3,306 TF- and miRNA-target transcripts were included for analysis. Note that each miRNA was required to express in at least 20 RNA Atlas profiled samples.

### Regulatory regions and regulator sequences

We predicted TF and lncRNA binding sites on 22,057 proximal promoters of 17,495 protein-coding genes. Each promoter is of length 2kbps, [+1kbp:-1kb] relative to the transcription start site. About one-third of protein-coding genes had multiple proximal promoters. The 3’-UTRs were used to predict miRNA or RBP binding sites. In total, we have compiled 38,183 3’-UTRs corresponding to 17495 protein-coding genes. The median length of all 3’-UTRs is 1004 bps. More than two-thirds of protein-coding genes had multiple 3’-UTRs.

While identifying lncRNAs act as miRNA, RBP, and TF decoys, we searched for binding sites of these regulators throughout the whole lncRNA transcript sequence. Similarly, we identified lncRNA binding sites in promoters that match any potential binding domains of lncRNAs without consideration to their potential structures. We applied triplexator^76^ v1.3.2 and TargetScan^77^ v6.0 to predict sequence-based triple-helix (or triplex) structures and miRNA binding sites, respectively. In total, we considered sequences of 11,799 mature miRNAs, 71,950 lncRNA transcripts (or 38,008 genes), and 33,956 circRNAs. We note that in subsequent analyses only RNA species that were expressed in at least 50% of samples were considered. Their sequences, including asRNAs, lincRNAs, and circRNAs, were extracted from the human genome assembly GRCh38 stored at the UCSC Genome Browser using twoBitToFa^42^.

### Prediction of TF-targets

We predicted targets for 641 human TFs based on both sequence and expression evidence. First, each predicted TF-target was required to have significant binding evidence from either 751 ENCODE ChIP-seq^30,78^ profiles or 1,631 human TF PWMs for 108 and 641 TFs, respectively. Second, we required each TF-target pair to exhibit significant co-expression pattern across RNA Atlas profiles samples.

ENCODE ChIP-seq data sets were profiled in 37 immortal cell lines and >60% of them are in K562 (n=121 for 61 TFs), GM12878 (n=113 for 64 TFs), HepG2 (n=97 for 51 TFs), A549 (n=67 for 27 TFs), and H1-hESC (n=62 for 36 TFs). More than one-third of TFs had at least 2 replicates in the same cell line. Human TF PWMs were collected from five sources including motifs annotated in Factorbook^79^ (see Table S2 in their paper; n=86 for 76 TFs), motifs of quality A-D in HOCOMOCO v9^80^ (n=429 for 397 TFs), high-confidence motifs in HumanTF^81^ (see Table S3 in their paper; n=659 for 361 TFs), JASPAR^82^ v5_alpha (n=104 for 100 TFs), and SwissRegulon^83^ downloaded on 03/18/2014 (n=353 for 333 TFs). To avoid matrix entries of value 0, a pseudo-count 1 was added to each entry before calculating the relative occurrence frequencies of nucleotides at each position.

We interrogated each of 22,057 proximal promoters to see if there is a significant ChIP-seq peak (Q-value<1E-10) or PWM-based binding site (P < 1×10^−5^). The significance of motif scores on either forward or reverse strand of the proximal promoters were compared to 5’-flanking regions of length 2kbps of their cognate proximal promoters using the CREAD^84,85^ package. Binding site evidence across multiple promoters associated with the same gene were aggregated to produce gene-level binding evidence. For any protein-coding gene that satisfied this sequence-based constraint, we further required significant distance correlation (dCor)^86^ at P < 1×10^−9^, as calculated using expression profiles of their regulating TFs and cognate protein-coding targets profiled in RNA Atlas. Note that only TFs and target genes of non-zero median absolute deviation (MAD) score were included for analysis. We applied permutation testing to estimate the significance of dCor by shuffling TF’s expression 100K times and then calculated the randomized dCor values. These values were used to fit parameters for a generalized extreme value (GEV) distribution using the MATLAB gevfit routine to obtain a non-parametric p-value lower than 1E-5 from the cumulative density of the resulting GEV distribution. For TF-targets passed both sequence and expression constraints were investigated for transcriptional lncRNA modulation. We predicted 115,565 TF-target genes significantly modulated by lncRNAs. Moreover, 160,227 TF-target interactions had target transcripts of adequate exonic and intronic coverage to compute m/p-ratio profiles.

### Prediction of miRNA- and RBP-targets

We predicted targets of both types of post-transcriptional regulators through a two-step approach by requiring both sequence- and expression-based evidence. Specifically, 3’-UTRs of protein-coding transcripts and whole lncRNA transcripts were scanned for miRNA binding sites conserved across species (context score < −0.2) by TargetScan^77^ v6.0 and significant RBP binding peaks at P < 1×10^−10^. ENCODE eCLIP^87^ datasets for 115 RBPs profiled in two human cancer cell lines, i.e., K562 and HepG2, were downloaded from UCSC Genome Browser. Among them, 66 and 49 RBPs were available in either one or two cell lines, respectively. Each RBP-cell line pair was performed in duplicates. Binding site evidence across multiple 3’ UTRs associated with the same gene were aggregated to produce gene-level binding evidence. We then asked if any pair of gene, either coding or non-coding, shared a significantly large common regulator program at adjusted pFET < 0.01. For each qualified gene pair and their common regulators, we measured if correlation changes between a common miRNA/RBP and any of these two genes had evidence for being modulated by lncRNA expressions using delta dCor; see the section “lncRNA target predictions using LongHorn.” below. A pair of regulator-target significantly modulated by at least one lncRNA at P < 0.05 was finally selected. miRNA/RBP-targets that passed both sequence and expression constraints were investigated for post-transcriptional lncRNA modulation. In total, 371,591 predicted miRNA-target interactions were identified as significantly modulated by lncRNAs and, among them, 33,602 miRNA-target transcripts had adequate exonic, intronic, and m/p-ratio reads and could be included in further analyses to compare correlations of regulator and target mRNA and pre-mRNA expression profiles. Note that, similar to experimentally verified miRNA targets, each miRNA, including both miRBase-annotated and RNA Atlas-identified miRNAs, was required to be expressed in at least 20 RNA Atlas profiled samples. RBPs were required to have a non-zero MAD score.

### Distance correlation

For each experimentally-verified and LongHorn-inferred TF- and miRNA-target, we applied distance correlation (dCor)^86^ to measure coexpression patterns between a regulator, namely, a TF, RBP, or miRNA, and its target using the target’s mRNA (exonic), pre-mRNA (intronic), or m/p-ratio profiles. Distance correlation is able to capture nonlinear relationship between two variables, which is a common scenario in the biological world. The dCor value is always non-negative and zero means that two variables are completely independent^86^. To better incorporate intrinsic noise of each type of expression, we randomly selected 10K pairs of TF- and miRNA-targets from all profiled genes without replacement, and then calculated their averaged dCor estimates (or baseline) using target’s mRNA, pre-mRNA, and m/p-ratio profiles. Dependent on the expression type used for computing the dCor between a regulator-target pair, the corresponding dCor baseline was taken from the computed dCor to become the adjusted dCor.

### lncRNA target predictions using LongHorn

LongHorn^25^ predicts lncRNA interactions through respectively integrating statistical evidence from modulation of transcriptional and post-transcriptional regulation by TFs, miRNAs, and RBPs. Transcriptional lncRNAs can physically interact with either proximal promoters (TR:Guide and TR:Co-factor) or TFs (TR:Guide and TR:Decoy) to alter their target pre-mRNA abundance. Guide and Co-factor lncRNAs form RNA-DNA triple-helix structures (triplex in short) with proximal promoters^76^. The former recruit TFs to bind their transcript’s promoter regions and synergistically enhance the transcription rates of their targets. The latter can either activate or inhibit the regulatory activities of TFs, which have their own binding sites on the same promoter. Decoy lncRNA, in turn, influences target pre-mRNA transcription and expression by altering the amount of TF and miRNA/RBP molecules available to target proximal promoters and 3’-UTRs. In our model, lncRNAs can regulate target pre-mRNA (TR:Decoy) and mRNA (PTR:Decoy) abundances in nucleus and cytoplasm. Note that, while predicting lncRNAs acting as guides or decoys of TFs, we used PWMs to scan TF binding sites from on lncRNA transcript sequences.

LongHorn reverse-engineers transcriptional and post-transcriptional interactions on a genome-wide basis at first; see above sections of “Prediction of TF-targets” and “Prediction of miRNA- and RBP-targets”. To estimate the significance of modulation, we calculated delta dCor for each triplet that is consist of an lncRNA, a regulator (TF/miRNA/RBP), and a protein-coding target. According to the lncRNA expression in each triplet, we partitioned RNA Atlas samples into four quartiles, from the lowest to the highest, and required this lncRNA to satisfy two constraints including 1) it was not correlated with the regulator (P > 0.1) (independence constraint) and 2) its expression fold change was > 2x between the fourth and the first quartiles of the samples (range constraint). Then, comparing the first and the fourth sample quartile, we required a nonparametric P < 0.05 for the delta dCor between the regulator and the target against a bootstrapping-based null hypothesis. For significant triplets that are associated with the same lncRNA-target pair, we integrated their p-values across their common regulators using either the Fisher’s method^88^ (transcriptional) or the weighted Brown’s method^89^ (post-transcriptional). While combining significance from multiple tests, the Brown’s method takes into account miRNAs and RBPs in the same genomic cluster, which are often co-expressed, to avoid inflating the integrated p-values. TargetScan context scores were used to sort predicted miRNA binding sites from lowest to highest; we then used their percentile ranks as weights for integrating p-values of significant triplets. After integration, we set a cutoff of adjusted p<0.01 for significant lncRNA-target pairs. Note that, both lncRNAs and their protein-coding targets were required to have a non-zero MAD score across profiled RNA Atlas samples.

### Stringent lncRNA-target set

We defined a stringent lncRNA-target set by evaluating the deviations of the adjusted dCor between each mediating regulator and the target pre-mRNA or m/p-ratio expression profiles. Specifically, LongHorn predicted lncRNA-target pairs by providing 1) target identities, 2) regulation model including transcriptional or post-transcriptional interactions, and 3) the list of regulators that are predicted to mediate the interactions. Each lncRNA-target interaction was associated with two distributions of the adjusted dCor: one was using pre-mRNA and the other using m/p-ratio expression profiles of the target. We anticipated that post-transcriptional targets of lncRNAs will have higher adjusted dCor values with their mediating regulators while using m/p-ratio than pre-mRNA expression profiles of targets. Yet, for transcriptional targets of lncRNAs, the relationships were expected to be in in the opposite direction—with m/p-ratio less correlated than the pre-mRNA expression profiles. We evaluated differences between these two distributions by the Student’s T test and required significance at p<0.05.

### Transcriptional and post-transcriptional specialists

To identify lncRNA specialists with unusual number of transcriptional or post-transcriptional interactions, we first normalize the size of LongHorn-inferred transcriptional and post-transcriptional interactomes to obtain a scaling ratio (σ). In RNA Atlas, LongHorn predicted 488,403 and 18,256 targets whose regulators’ activities were transcriptionally and post-transcriptionally modulated by lncRNA, respectively. Namely, the scaling ratio is 26.753 for transcriptional interactions. For each RNA Atlas profiled lncRNA, including asRNAs, lincRNAs, and circRNAs, with at least 10 predicted targets, we calculated the adjusted fold change (adjFC) using the following formula to determine if it is a specialist:

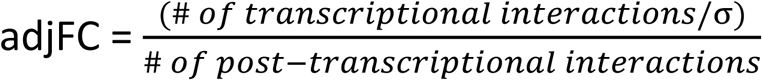

The lncRNA is a transcriptional or post-transcriptional specialist if the adjFC is larger than 2 or smaller than 0.5, respectively. By calculating this statistic, we revealed lncRNAs extensively involved in pathways of transcriptional or post-transcriptional gene regulation.

### Hallmark Gene Set enrichment analysis

We sought to study if lncRNAs regulate key biological pathways through searching for significant overlaps between their LongHorn-inferred targets and 50 MSigDB Hallmark Gene Sets^34^, which can be broken into 8 basic categories including Cellular Component, Development, DNA Damage, Immune, Metabolic, Proliferation, and Signaling Pathways. We calculated p values for the significance of overlap using Fisher’s Exact Test and adjusted for multiple comparisons based on Bonferroni correction. For each lncRNA-gene set pair, an adjusted p-value lower than 0.01 was considered to be a significant association.

## Data availability

All types of RNA entities can be readily explored via the online R2: Genomics analysis and visualization platform (http://r2.amc.nl), and via a dedicated accessible portal (http://r2platform.com/rna_atlas). This portal includes Genome browser profiles for the total RNA as well as polyA tracks for all samples. All samples can also be used for correlations, differetntial signals and many more analyses. In addition, the LongHorn results, described in this manuscript can be explored.

The raw data (fastq files) and processed expression measurement tables from all RNA biotypes across samples have been deposited in NCBI’s Gene Expression Omnibus^90^ and are accessible through GEO Series accession number GSE138734 (ncbi.nlm.nih.gov/geo/query/acc.cgi?acc=GSE138734).

## Author contributions

P.M., J.V. and P.S. conceived the idea, designed and supervised the project. L.R. and H.-S.C. contributed to the implementation and design of most bioinformatic analyses. H.-S.C., T. - W.C. and P.S. performed the analyses related to regulatory interactions of non-coding RNAs. F.A.C. and K.D.P. performed the analyses to select a stringent set of genes and contributed to quality validation of the transcriptome. S.G., S.K. and G.P.S. generated and sequenced the polyA and total RNA libraries. P.-J.V. performed the evaluation of coding potential and analyses of mass spectrometry data. R.C. and Y. S. contributed to the analyses of RNA biotype expression and sample ontology associations. J.N. performed the polyA-minus sequencing and the qPCR experiments. K.V. and J.N. generated and sequenced the small RNA libraries,. J.A. implemented the identification of novel miRNAs and sequence motifs analysis. S.L. designed the primers for the qPCR experiments. T.G. and T.D.M. performed the imprinting analyses. T.B.H. and J. Kjems implemented the circRNA identification workflow. N.N. developed the polyA-minus sequencing protocol. T.T., K.V. and K.R.B. provided immune system-related cell lines and cell types. N.D., G.A., M.R.W. and A.U. performed analyses and annotation of circRNAs. J. Koster developed dedicated tools to analyze RNA atlas data and results and implemented them in the online portal R2. P.M. led the writing of the manuscript in collaboration with L.R., H.-S.C. and P.S. L.R., H.-S.C., G.P.S., J.V., P.S. and P.M. contributed to the conceptualization, interpretation and discussion of results. All authors commented on the manuscript and contributed to the presentation of the data and results.

## Acknowledgements

F.A.C. is supported by Special Research Fund (BOF) scholarship of Ghent University (BOF.DOC.2017.0026.01). A.U. is supported by research funding from the NHMRC (Australia) and Leukemia & Lymphoma Society, Leukemia Foundation and Snowdome Foundation. G.A. is supported by a postgraduate scholarship from the Translational Cancer Research Network. M.R.W. and N.D. acknowledge support from the Australian Government NCRIS program, administered by Bioplatforms Australia. We thank Angelica Barr, Smita Pathak, Lisa Way, Anthony Mai for their contribution in library preparation, and Allison Yunghans, Erich Jaeger and Ali Moshrefi for their assistance in library organization and sequencing/tracking/data management. This project was funded by the European Union’s Horizon 2020 research and innovation programme under grant agreements 668858 and 826121 to P.M., P.S., and J.Koster.

## Supplemental Tables

Table S1. RNA atlas samples.

Table S2. Annotation of filtered known and novel mRNA, lincRNA and asRNA genes assembled from all polyA and total RNA libraries.

Table S3. Annotation of known and novel filtered circRNAs identified across all total RNA libraries.

Table S4. Annotation of known and novel filtered mature miRNAs identified across all small RNA libraries.

Table S5. Annotation of known and novel filtered miRNAs precursors identified across all small RNA libraries.

Table S6. Novel candidate mRNAs with matching peptides from mass spectrometry data in the Human Proteome Map.

Table S7. Table of known polyadenylated and non-polyadenylated genes used as reference for the polyadenylation status classification. This list was generated based on Yang et al. (2011).

Table S8. List and annotation of 160 genes with variable polyadenylation status across samples.

Table S9. Overview of significantly imprinted genes in 203 tissue and cell type samples.

Table S10-12. mRNA, pre-mRNA, and m/p-ratio expression profiles of protein coding transcripts (quantile-normalized and log2-transformed). (Related to Figure 5)

Table S13. Experimentally-verified targets of canonical regulators including TFs and miRNAs. (Related to Figure 5)

Table S14. LongHorn-inferred targets of canonical regulators including TFs and miRNAs. (Related to Figure 5)

Table S15. The stringent set of miRBase and RNA-Atlas miRNAs and their LongHorn-inferred targets. (Related to Figure 5)

Table S16. lncRNA network predicted by LongHorn using RNA-Atlas expression profiles. (Related to Figure 6)

Table S17. Significance of correlation deviations for LongHorn-inferred lncRNA targets. (Related to Figure 6)

Table S18. Significant lncRNAs in Hallmark Gene Set enrichment analysis. (Related to Figure 6)

Table S19. qPCR validation experiment with 110 single-exon genes.

Table S20. Annotation of fusion genes labels retrieved by FusionCatcher and criteria used for filtering false positives.

