## Supplementary material for "The RNA Atlas, a single nucleotide resolution map of the human transcriptome": Figure S

Figure S1

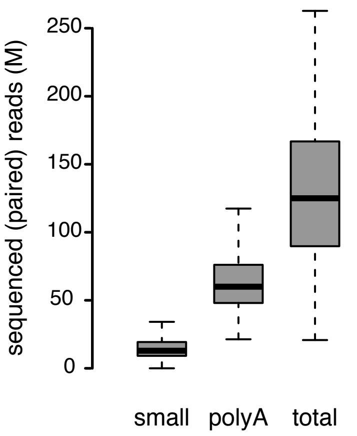


Sequenced read counts per library type. Distribution of total sequenced reads (small RNA-seq) or read pairs (polyA and total RNA-seq) across samples for the different data sets.

Figure S2

A

B

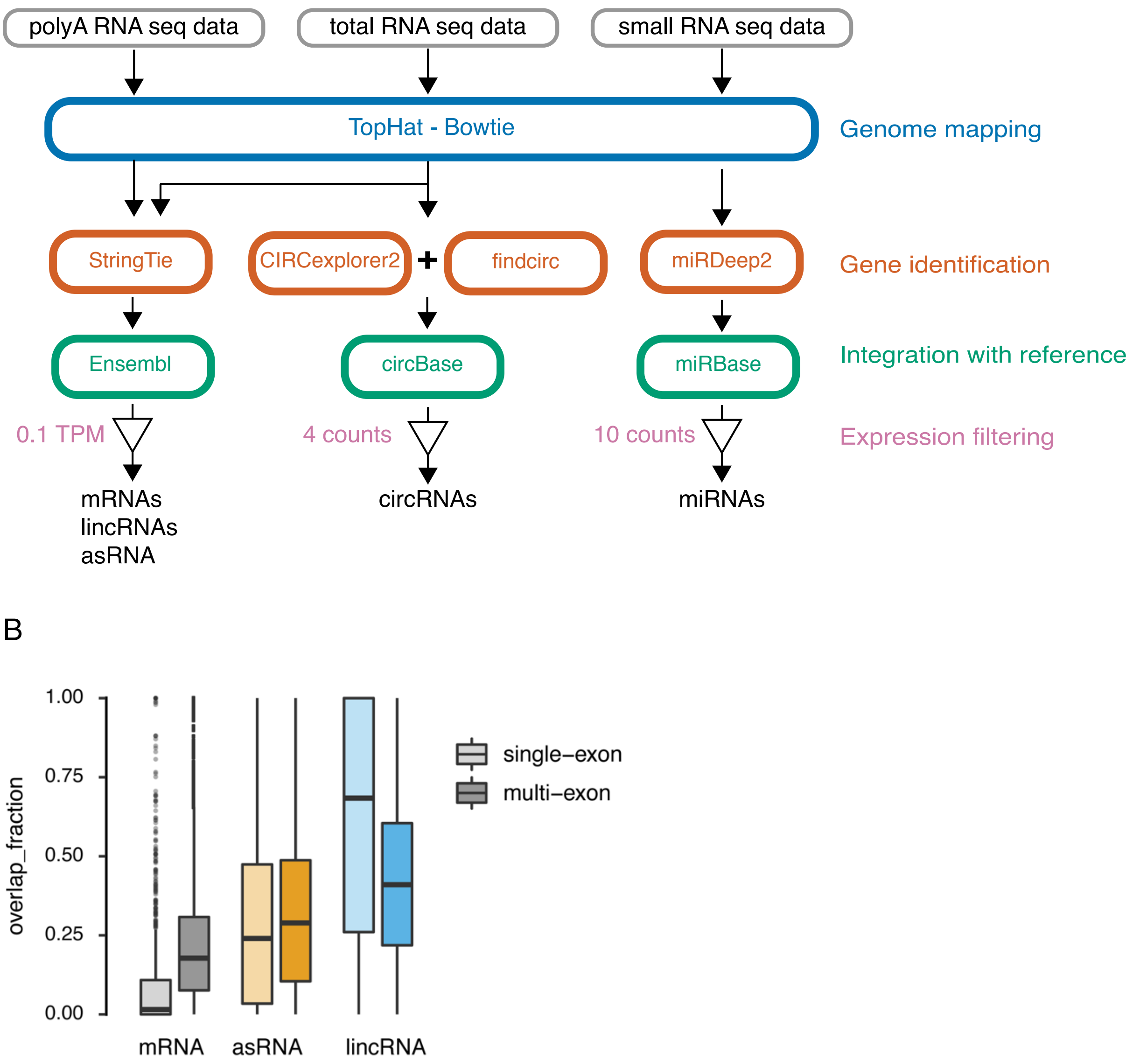


(A) Overview of the workflow used for assembly of the RNA atlas transcriptome. After expression filtering of mRNAs, lincRNAs and asRNAs, overlap with repetitive elements was calculated for further filtering as described in the methods section. (B) Distribution of the fraction of exon sequence overlap with repeats. For each RNA biotype, the left and right boxes correspond to exons from single-exon and multi-exon genes, respectively.

Figure S3

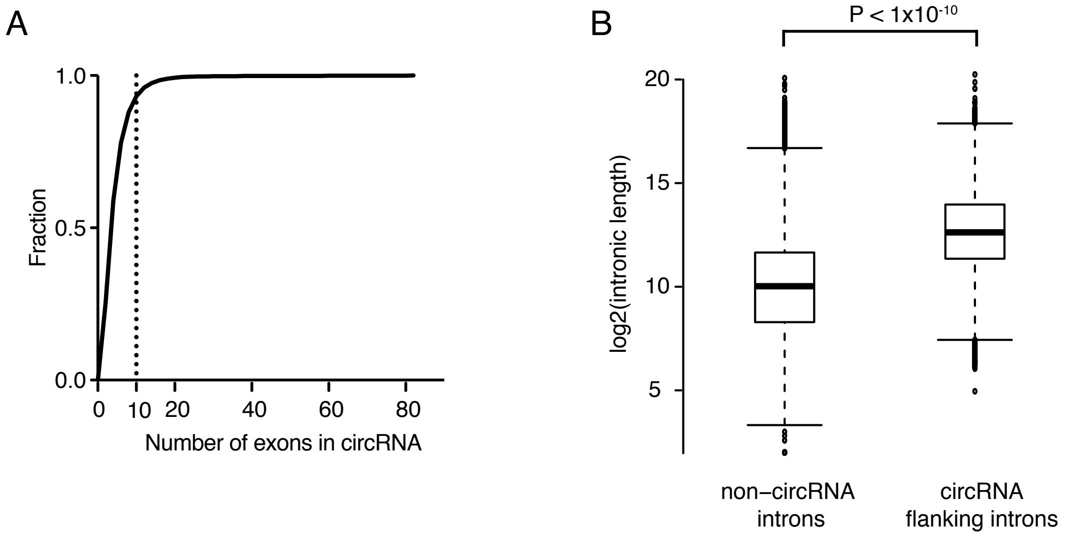


Properties of circRNAs. (A) Cumulative fraction of number of exons per circRNA. (B) Intron length distributions for introns flanking circRNAs and introns not flanking any circRNA.

Figure S4

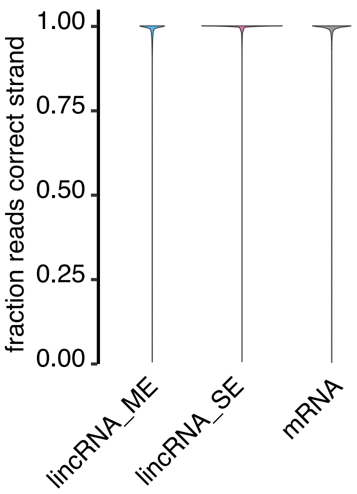


Fraction of reads mapping to the correct genomic strand in the sample with maximum expression for exons from multi- and single-exon lincRNAs (lincRNA_ME, and lincRNA_SE, respectively) and exons from mRNAs.

Figure S5

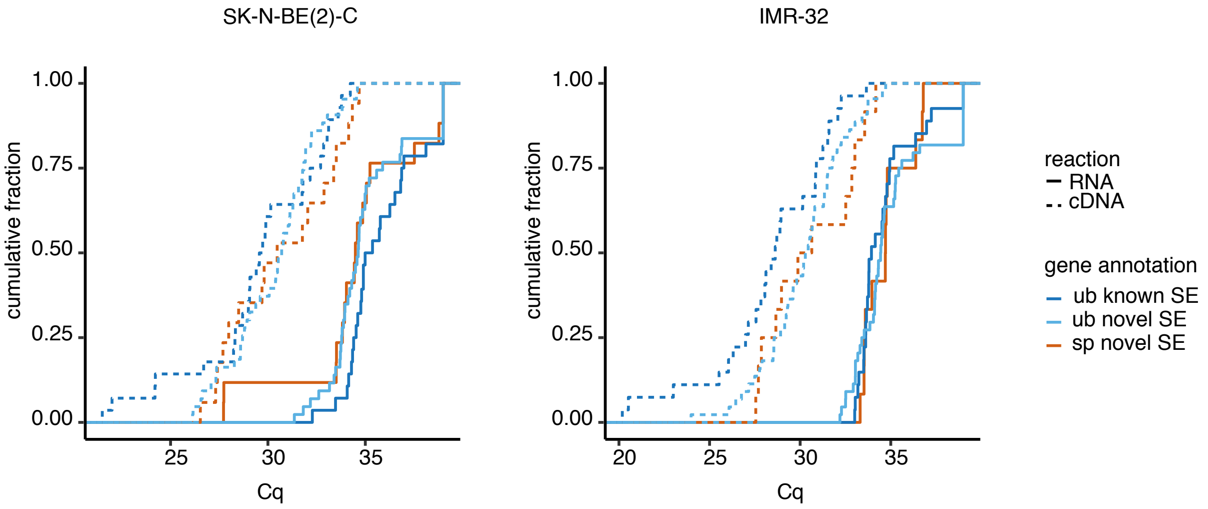


qPCR validation of novel single-exon genes. For each of the cell lines used, two qPCR reactions were performed, one using total RNA, and one using cDNA as template. Cumulative fractions of Cq-values obtained for each reaction are shown separately for three different sets of genes: ubiquitous known single-exon genes (ub known SE, dark blue), ubiquitous novel single-exon genes (ub novel SE, light blue) and novel single-exon genes specifically expressed in each cell line (sp novel SE, orange).

Figure S6

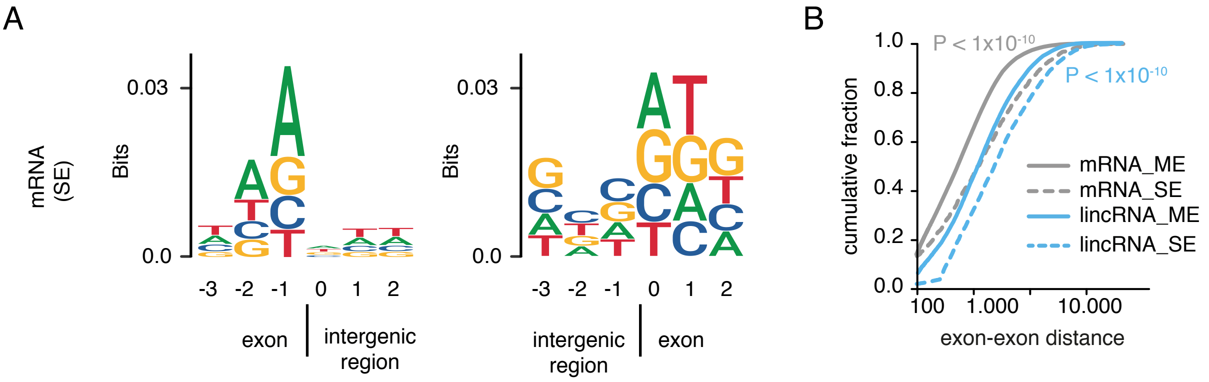


Expected properties of single exon genes. (A) Nucleotide frequencies at intergenic/exonic boundaries for single-exon mRNAs. (B) Cumulative fraction of the distance to the closest exon for multi-exon and single-exon mRNA exons (grey-colored lines, mRNA_ME and mRNA_SE, respectively), and multi-exon and single-exon lincRNA exons (light blue-colored lines, lincRNA_ME and lincRNA_SE, respectively). p-values were calculated using the Wilcoxon signed-rank test.

#### Figure S7

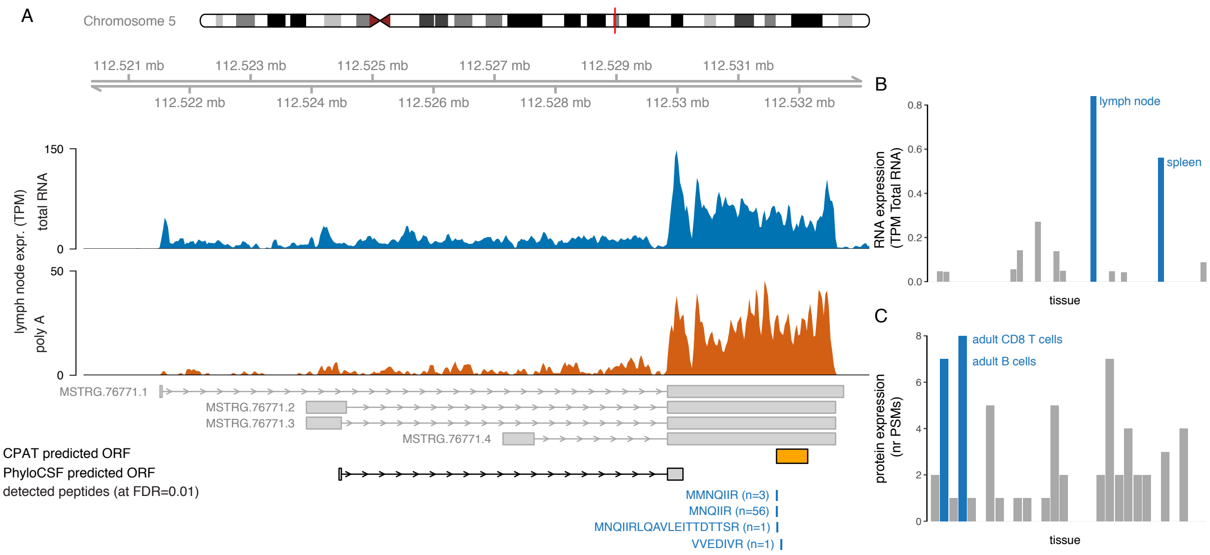


Example of a novel mRNA with matching peptides from public mass spectrometry data. (A) The different tracks show genomic coordinates of the identified gene, coverage profiles from total and polyA RNA-seq, transcript structure of the 4 assembled isoforms, predicted ORFs by CPAT and PhyloCSF and detected peptides matching the CPAT predicted ORF. Expression of this gene is enriched in immune related samples (blue bars) both at RNA (B, expression for isoform 3 is shown) and peptide (C) level.

#### Figure S8

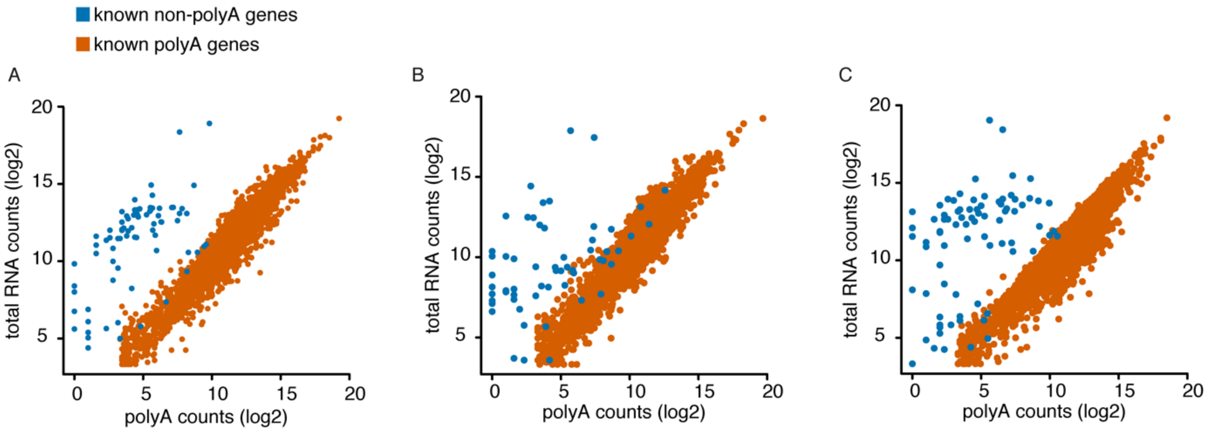


Correlation between log2 counts from total RNA-sequencing and polyA RNA-sequencing for known polyadenylated (orange) and non-polyadenylated (blue) genes in a cell type (A: human umbilical vein endothelial cell), a tissue (B: distal colon), and a cell line (C: K562).

Figure S9

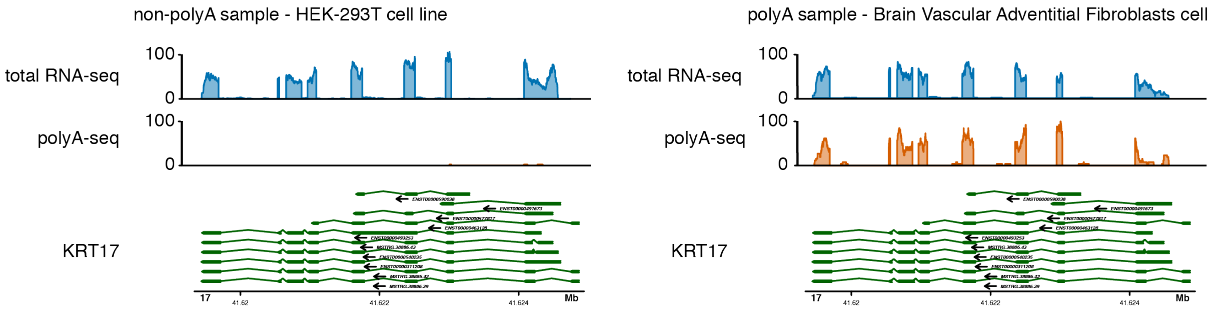


Example of a gene (KRT17) with variable polyadenylation across samples with no clear differences between the expression patterns at transcript level for polyadenylated and non-polyadenylated samples. Coverage profiles from total RNA-sequencing and polyA-sequencing are shown for a non-polyadenylated sample (left) and a polyadenylated sample (right).

Figure S10

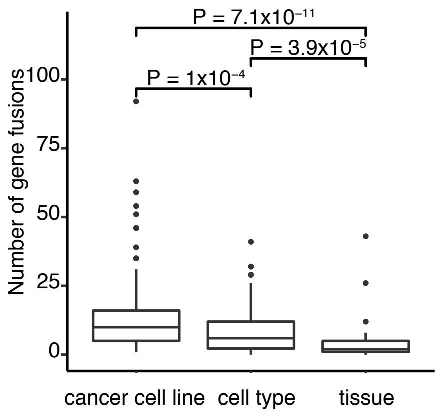


Distribution of the number of predicted fusion genes identified per sample, for different sample types. The significance of differences in number of fusion genes between sample types was assessed with the Wilcoxon signed-rank test.

Figure S11

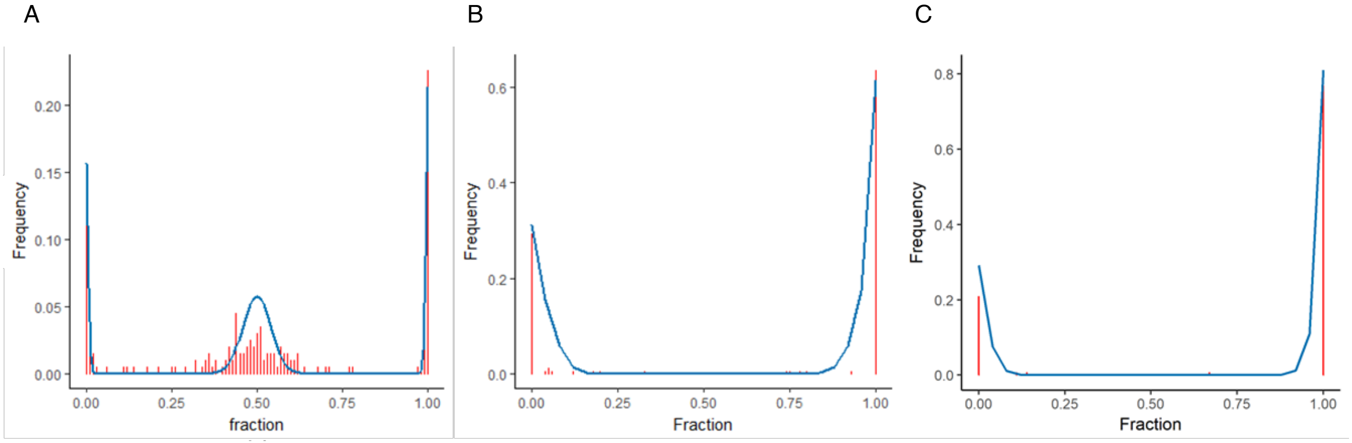


Imprinting analyses: Observed (red) and expected (blue) frequencies of alternative alleles for two SNP positions. (A) An example of a non-imprinted SNP in FAM20C (rs36138803, degree of imprinting (i) = 0). A clear heterozygous peak is present, i.e. a set of samples with an alternative allele fraction of roughly 50%. (B) A significantly imprinted SNP in PLAGL1 (rs17073273, adj. p-value = $9.2 \times 10^{-49}$ and i = 0.97). Here the heterozygous peak is eliminated and virtually no heterozygous samples are observed. (C) A significantly imprinted SNP in MIR381HG (rs35844276, adj. p-value = $7.5 \times 10^{-34}$ and i = 0.99).

Figure S12

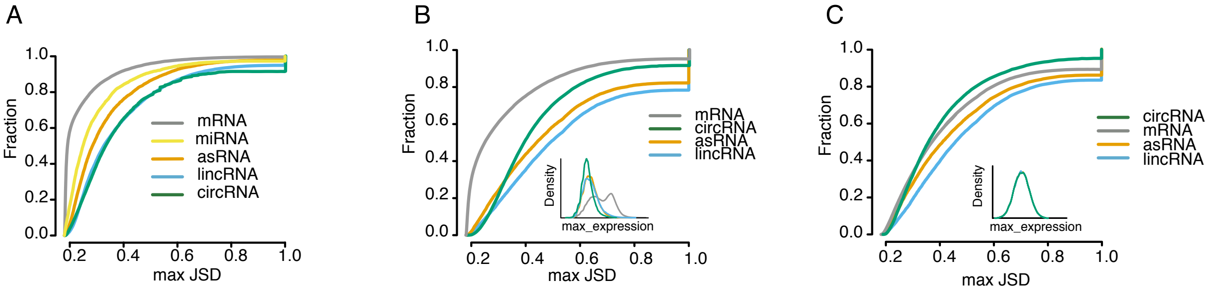


Specificity before and after correcting for expression distributions. In all cases, the specificity score for a given feature was calculated as the maximum Jensen-Shannon divergence. (A) Cumulative fraction of the specificity score at gene level for all RNA biotypes without correcting for differences in abundance. (B) Cumulative fraction of the specificity scores for all forward-spliced junctions from mRNAs, lincRNAs and asRNAs, and back-spliced junctions from circRNAs. The inset plot shows the density distributions for the maximum expression across samples for the different biotypes. The expression was calculated as junction counts scaled by library size. (C) Similar to (B), but correcting for differences in abundance distribution by performing a directed subsampling of junctions from the different biotypes to match a common expression distribution, as shown in the inset plot.

Figure S13

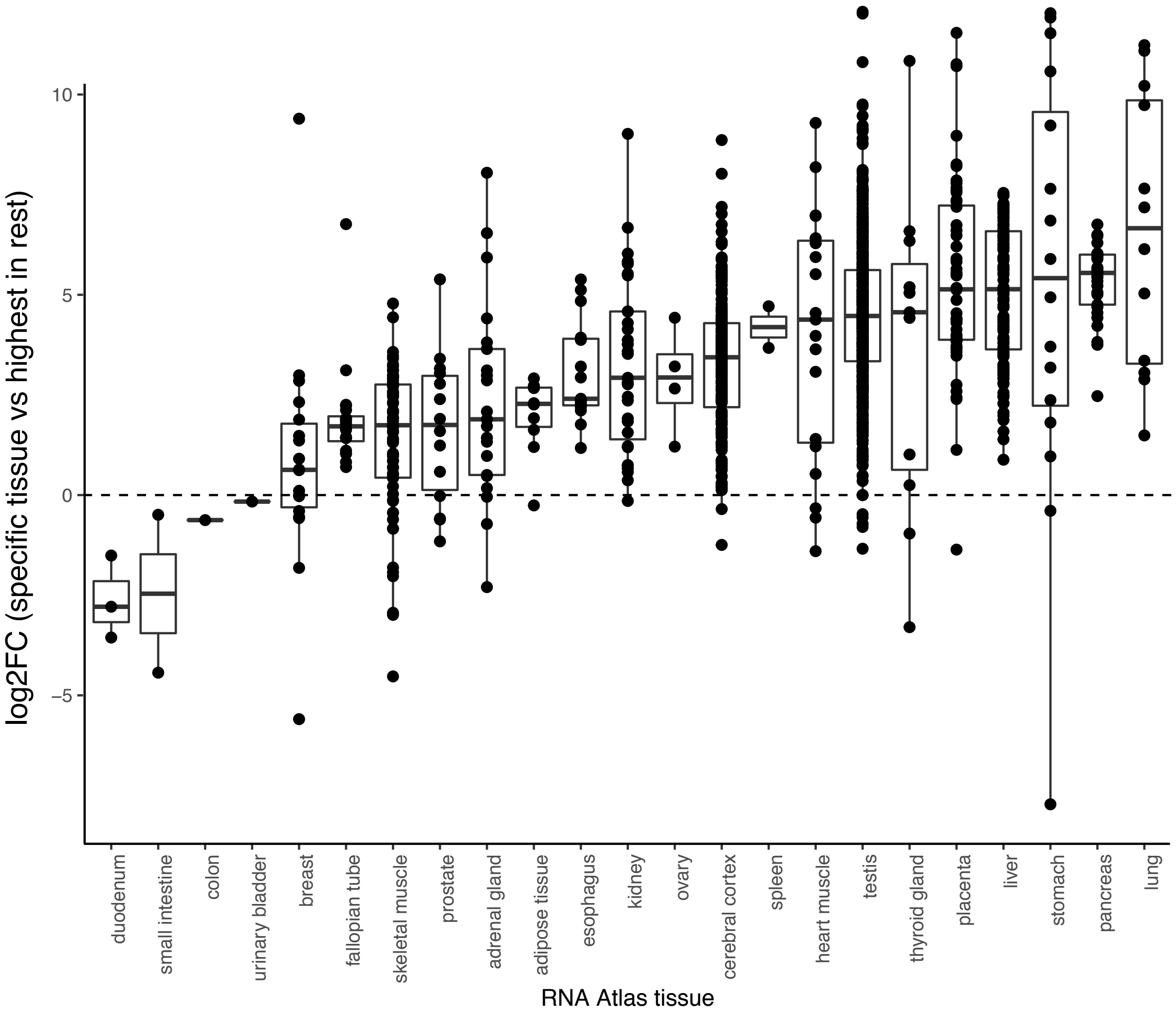


Cross validation of tissue-specific markers selected from the Human Protein Atlas, as explained in Methods. The y-axis shows the log2 fold-change between the expression of the tissue-specific marker in the matching RNA atlas tissue and its highest expression among the remaining 22 tissues.

### Figure S14

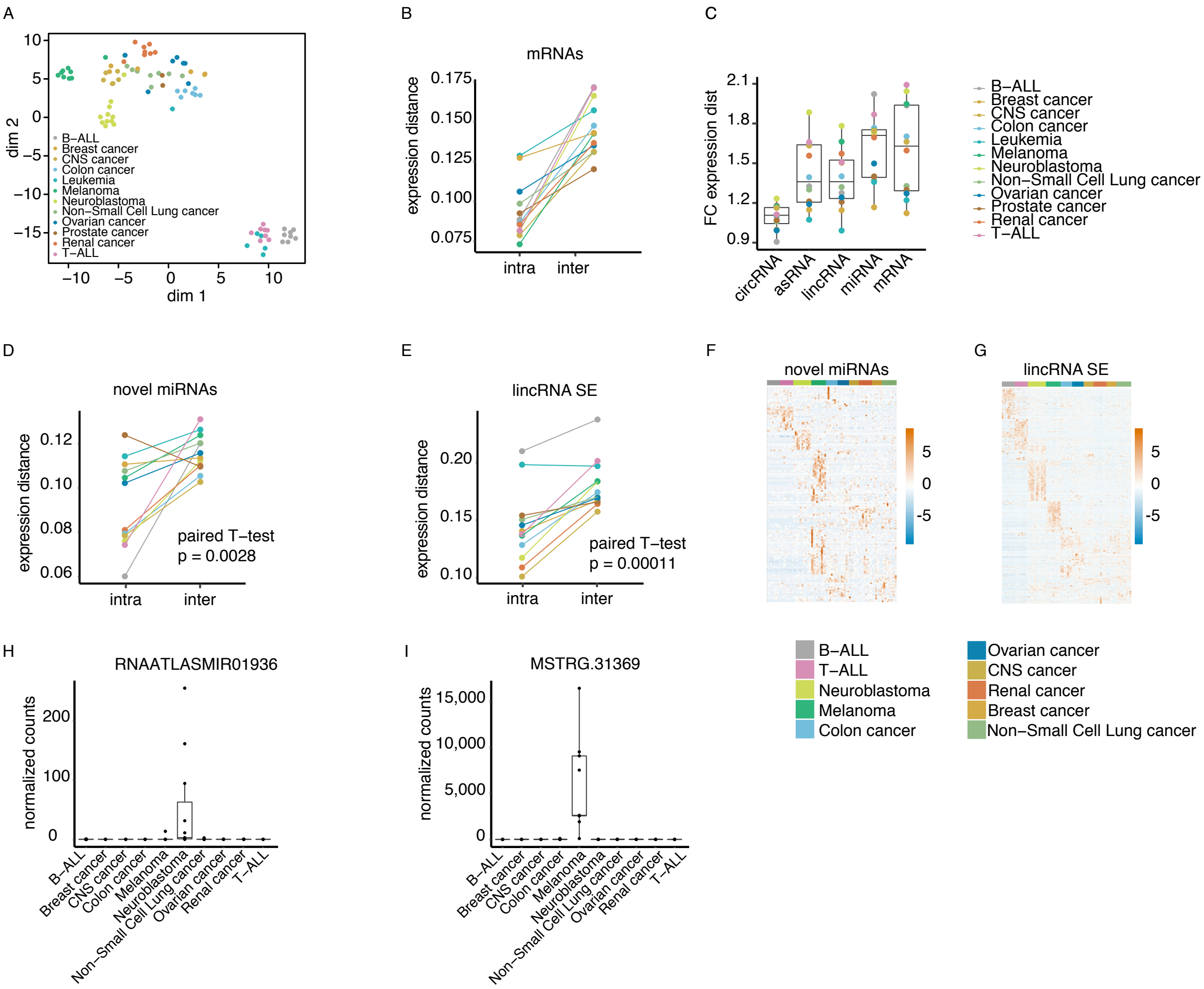


Association between cancer type and expression-distance within the cancer cell lines dataset. (A) t-SNE plot of the RNA atlas cancer cell lines based on mRNA expression. Samples are colored according to the cancer type. (B) For each cancer type, the median mRNA expression-based distances between pairs of samples within a cancer type(intra-distances) and between each cell line from the cancer type and all other cancer cell lines (inter-distances) are shown. (C) Distribution of fold changes between median inter- and intra-distances calculated based on expression of the different RNA biotypes. (D) Median intra- and inter-distances based on expression of novel miRNAs. (E) Median intra- and inter-distances based on expression of single-exon lincRNAs. (F) Expression heatmap for novel miRNAs significantly upregulated in each of the cancer types. (G) Expression heatmap for single-exon lincRNAs significantly upregulated in each of the cancer types. (H) Example of a neuroblastoma-specific novel miRNA. (I) Example of a melanoma-specific single-exon lincRNA.

#### Figure S15

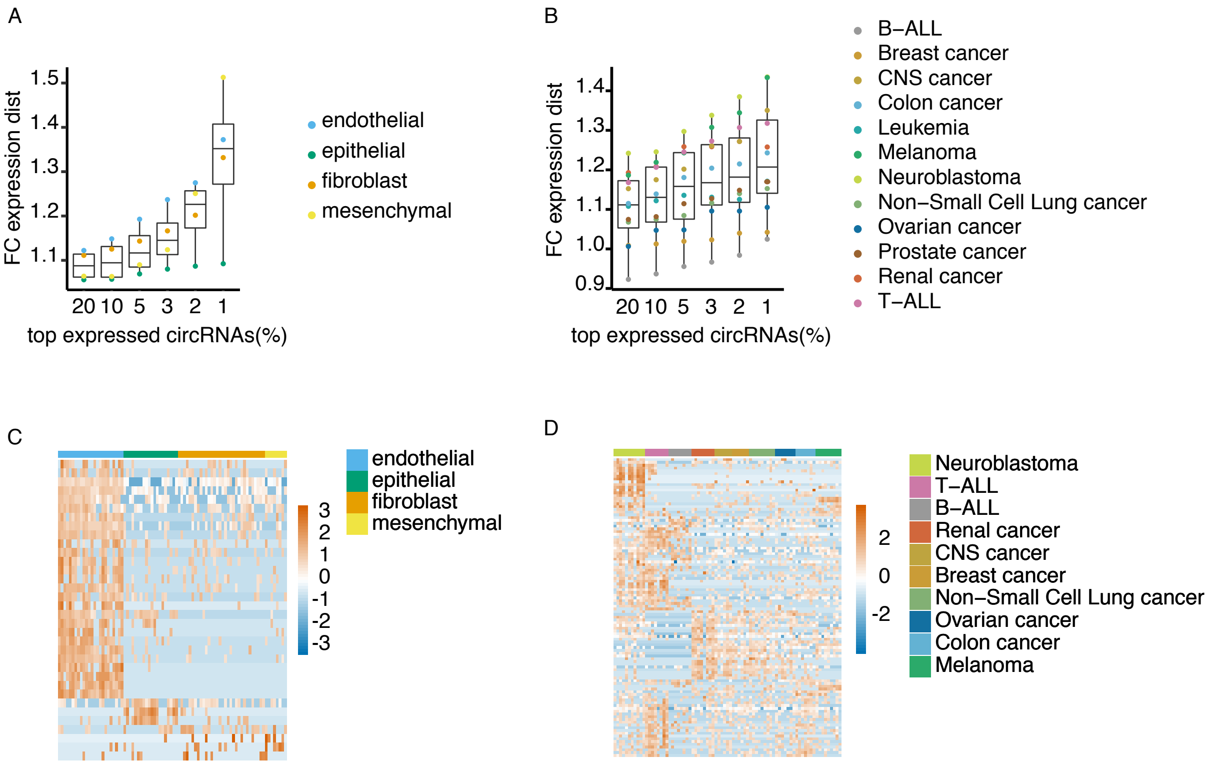


Analyses of circRNA expression distances between samples within and between biological subtypes. (A) Impact of selecting subsets of abundant circRNAs across samples on the fold change in expression distances within and between cell subtypes. (B) Impact of selecting subsets of abundant circRNAs across samples on the fold change in expression distances within and between cancer types. (C) Heatmap of identified differentially expressed circRNAs for each of the 4 subtypes studied. (D) Heatmap of identified differentially expressed circRNAs for each of the cancer types.

#### Figure S16

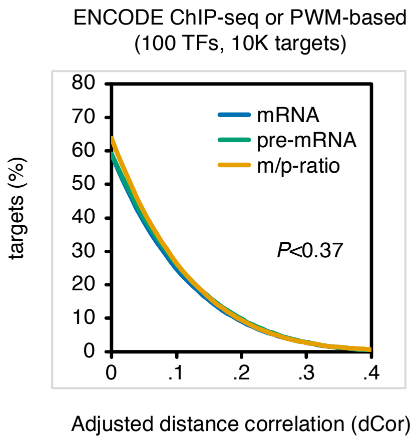


A list of 10K TF-target predictions with no supporting evidence from expression were randomly drawn and they did not show correlation differences between TF-target pre-mRNA and Ratio.

Figure S17

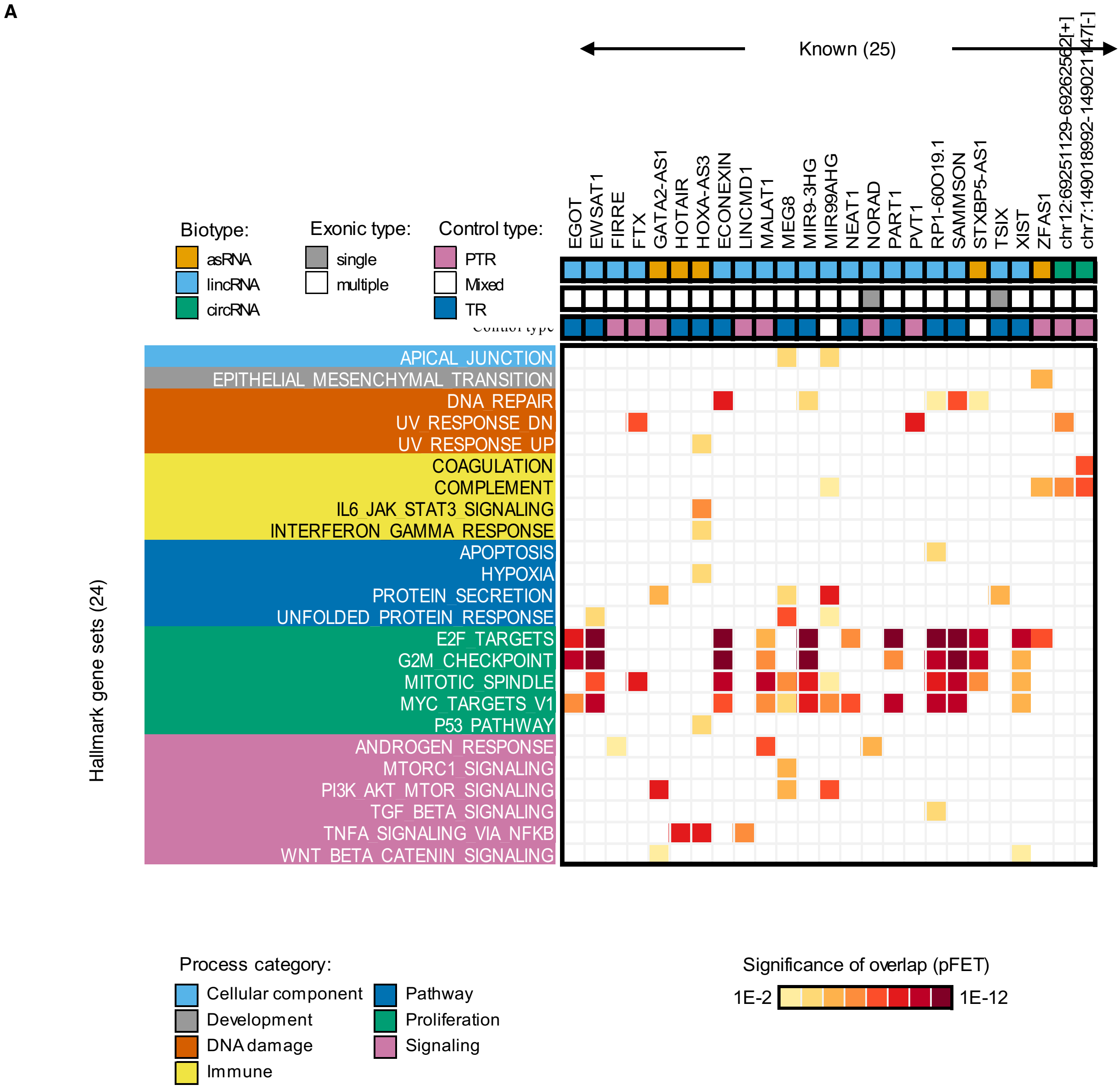
A
Known (25)
EGOT
EWSAT1
FIRRE
FTX
GATA2-AS1
HOTAIR
HOXA-AS3
ECONEXIN
LINCMD1
MALAT1
MEG8
MIR9-3HG
MIR99AHG
NEAT1
NORAD
PART1
PVT1
RP1-60O19.1
SAMMSON
STXBP5-AS1
TSIX
XIST
ZFAS1
chr12:69251129-69262562[+]
chr7:149018992-149021147[-]
Biotype:
asRNA
lincRNA
circRNA
Exonic type:
single
multiple
Control type:
PTR
Mixed
TR
Hallmark gene sets (24)
APICAL_JUNCTION
EPITHELIAL_MESENCHYMAL_TRANSITION
DNA_REPAIR
UV_RESPONSE_DN
UV_RESPONSE_UP
COAGULATION
COMPLEMENT
IL6_JAK_STAT3_SIGNALING
INTERFERON_GAMMA_RESPONSE
APOPTOSIS
HYPOXIA
PROTEIN_SECRETION
UNFOLDED_PROTEIN_RESPONSE
E2F_TARGETS
G2M_CHECKPOINT
MITOTIC_SPINDLE
MYC_TARGETS_V1
P53_PATHWAY
ANDROGEN_RESPONSE
MTORC1_SIGNALING
PI3K_AKT_MTOR_SIGNALING
TGF_BETA_SIGNALING
TNFA_SIGNALING_VIA_NFKB
WNT_BETA_CATENIN_SIGNALING
Process category:
Cellular component
Development
DNA damage
Immune
Pathway
Proliferation
Signaling
Significance of overlap (pFET)
1E-2
1E-12

B

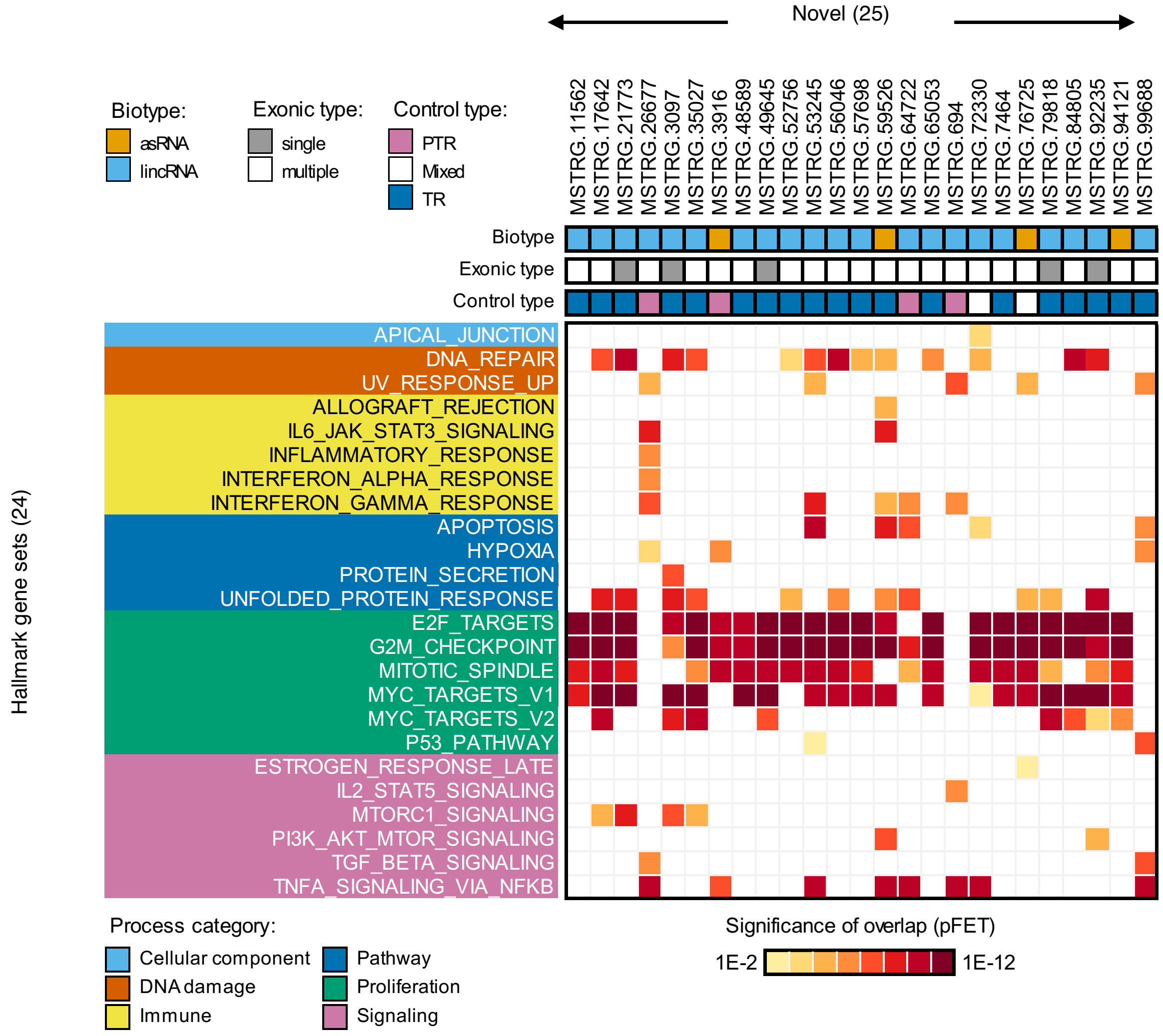


Enrichments for the predicted targets of twenty-five (A) previously cataloged (known) and (B) RNA Atlas (novel) lncRNAs in MsigDB hallmark pathways. lncRNAs are cataloged based on biotype, whether they are single- and multi-exon, and regulatory modality.
